# DNA repair functions are essential for bacterial stress defense but render antibiotic tolerant sub-populations susceptible to ciprofloxacin and gentamicin upon prolonged treatment

**DOI:** 10.1101/2025.02.03.636342

**Authors:** Yingkun Wan, Jiaqi Zheng, Heng Heng, Yang Tang, Lianwei Ye, Edward Wai-Chi Chan, Sheng Chen

## Abstract

The unresponsiveness of bacterial tolerant subpopulations to antibiotics has been attributed to physiological dormancy triggered by environmental stresses. In this work, we showed that protein and DNA synthesis activities in tolerant subpopulations that formed during nutrient starvation remained at a high level and only dropped gradually during the course of six days-starvation. Interestingly, upon decline of these activities, the tolerant subpopulations became susceptible to antibiotics that target protein and DNA synthesis, which was found to be required to support DNA repair functions essential for prolonged survival of the tolerant subpopulation. The increasing susceptibility of the bacterial tolerant subpopulation to antibiotics was also due to diminishing energy production, antioxidant defense and efflux functions, which resulted in weakening defense and further inhibition of synthesis and functions of DNA repair and other defense proteins. These findings confirm that bacterial tolerant subpopulations remain physiological active, and open new opportunities for combating bacterial tolerance with existing antibiotics.

## Introduction

The concept or definition of antibiotic or stress tolerance in bacteria has been vague among the scientific and medical fields ever since this phenomenon was discovered by Bigger in 1944 (Bigger, 1944). For decades, unresponsiveness of bacteria to antimicrobial treatment has been attributed to the fact that bacteria become physiologically dormant upon encountering environmental stresses such as nutrient starvation. The clinical importance of this phenomenon has not been recognized until recent years, when a number of studies began to show that bacterial antibiotic tolerance was responsible for causing chronic and recurrent infections, and predisposing development of antibiotic resistance (Zheng et al., 2022). It is therefore necessary to differentiate between antibiotic tolerance and resistance and delineate the physiological mechanisms that support maintenance of the tolerance phenotype. Antibiotic resistant organisms exhibit mechanisms that help withstand the antimicrobial effects of antibiotics and are able of replicating in the presence of antibiotics, whereas antibiotic tolerant bacterial sub-population refer to those which temporarily do not respond to antimicrobial treatment under specific environmental conditions such as nutrient starvation, even though the organism concerned is susceptible to antibiotics when it grows under favorable conditions. In other words, switching to a tolerance mode enables bacteria to survive against multiple stresses, including the antimicrobial effects of antibiotics. When the threat subsides and conditions become favorable, members of such tolerant population, which are commonly known as tolerant subpopulations, can rapidly resume growth. Antibiotic tolerant bacterial sub-population may pose even more serious threat to human health than antibiotic resistant strains, considering that such tolerant cells exist in all bacterial populations and that hostile conditions in the human body always render a certain proportion of any bacterial population tolerant to multiple antibiotics(Fung et al., 2010).

Various previous studies showed that bacteria actively switched off the key physiological functions and metabolic activities under unfavorable conditions (Wood et al., 2013). Previous works that aimed at delineating bacterial tolerance mechanisms therefore focused on stress response regulated by the alarmone (p)ppGpp and the toxin-antitoxin system, which mainly mediates shutdown of physiological activities (Harms et al., 2016). Nevertheless, when bacteria cease to replicate, cell wall and DNA synthesis activities are expected to ground to a halt, rendering β-lactam and fluoroquinolone antibiotics unable to exert their antimicrobial effect. It is also likely that protein synthesis activities decrease substantially when bacteria become physiological dormant, hence the stress tolerant sub-population should also be tolerant to the aminoglycosides.

A number of recent studies, however, indicated that bacteria do not simply turn off all physiological functions in order to exhibit tolerance to antibiotic or environmental stresses. In 2016, Pu et al showed that bacterial tolerant sub-population exhibited a wide range of efflux activities, which presumably played a role in exuding toxic metabolites out of the cell to enhance survival fitness (Pu et al., 2016). In fact, some of the earlier studies had reported up-regulated efflux and other physiological functions in bacteria that encountered environmental stresses. Adams et al detected efflux activity in bacteria that exhibited phenotypic stress tolerance induced by macrophages (Adams et al., 2011). Tolerant bacteria cells were also found elicit active stress responses such as INH-activation (Wakamoto et al., 2013), energy production (Ma et al., 2010), the stringent response which involves the synthesis of the alarmone ppGpp (Bokinsky et al., 2013), the active σ^S^ stress response (Radzikowski et al., 2016), virulence gene expression (Arnoldini et al., 2014), SOS response induction (Goormaghtigh & Van Melderen, 2019) and fine-tuning of metabolic and respiratory functions (Orman & Brynildsen, 2015).

More recently, our laboratory also discovered that antibiotic tolerant subpopulations can actively maintain the transmembrane proton motive force (PMF) and specific efflux functions, and that the expression level of a number of putative tolerance genes recorded during nutrient starvation, a condition known to trigger onset of antibiotic tolerance, was much higher than that recorded in exponentially growing cells (Wang et al., 2021). We found that the ability to generate and maintain PMF was important for maintaining the antibiotic tolerance phenotype in bacteria subjected to prolonged starvation; hence abolishment of such ability by deletion of the PMF maintenance gene *pspA* and genes that encode the electron transport chain components, such as *ndh* and *nuoI*, resulted in increased susceptibility of the tolerant subpopulations of *E. coli* to ampicillin as a result of diminishing efflux activity and the associated increased accumulation of antibiotics in the intracellular compartment of bacterial cells. Further studies showed that a number of genes that encode efflux pumps, membrane transporters and components of electron transport chain were involved in maintenance of tolerant bacteria status (Wan et al., 2023; Wan et al., 2021). This observation is intriguing, as tolerant subpopulations that form during nutrient starvation are supposed to be physiologically inert and not able to replicate; hence accumulation of antibiotics in the bacterial cells should not exhibit any inhibitory effect on cell wall synthesis activity. We hypothesize that bacterial tolerant subpopulations are actually physiologically and metabolically active so that they should remain susceptible to antibiotics, and that the reason why a bacterial subpopulation is tolerant to antibiotics is because they also undergo active efflux to pump antibiotics out of the cells. When the ability to maintain efflux functions weakens, the tolerant bacterial cells would become increasingly susceptible to antibiotics if their cellular activities persist. In this work, we tested this hypothesis and investigated the cellular activities that occur in tolerant cells that form during nutrient starvation by testing various gene deletion mutants and using different types of antibiotics to probe the physiological status of the tolerant cells. Our findings confirm that antibiotic tolerant subpopulations have to actively undergo cell wall, protein and DNA synthesis even upon encountering nutrient starvation for a prolonged period in order to maintain the tolerance status. When efflux and other defense functions in such subpopulations weakened during prolonged starvation, the survival fitness of tolerant subpopulations would be compromised. Lower efflux would also lead to accumulation of antibiotics which would inhibit the essential tolerance-maintaining functions that remain switched on in the tolerant cells, rendering them susceptible to multiple antibiotics. These observations have important implications in future development of novel strategies to combat bacterial antibiotic tolerance.

## Results

This work aimed at presenting an overview of the physiological changes in bacterial populations that develop tolerance to multiple antibiotics upon encountering adverse environmental conditions. We first tested the changes in antibiotic tolerance levels of nine strains of different species and varied antibiotic susceptibility status over a period of six days when nutrients were completely deprived of. We chose six days as the study period as this is a typical time course for antimicrobial treatment. The test strains included *Escherichia coli* BW25113, SZ457-1, *Salmonella* PY1, SR5, *Klebsiella pneumoniae* PM15, EH62, *Acinetobacter baumannii* ATCC17978, CPC35 and *Pseudomonas aeruginosa* PAER00606, PAO1. Among them, SZ457-1, PM15, EH62, SR5, PAER00606, PAO1, ATCC17978, CPC35 exhibited varied degrees of reduced susceptibility to different antibiotics (Table S1). Another purpose of this experiment was to investigate whether different bacterial species exhibited differential ability to maintain a tolerance phenotype for a prolonged period, and whether antibiotic sensitive and resistant strains exhibited different levels of antibiotic tolerance when encountering starvation stress.

### Bacterial tolerant subpopulations become increasingly susceptible to ciprofloxacin and gentamicin upon prolonged starvation

As shown in Figure 1 and S1, ten times MIC of ampicillin, gentamicin and ciprofloxacin could not eradicate the organisms in the first 24 hours, indicating that a large proportion of the bacterial population had become antibiotic tolerant under starvation stress. As tolerant subpopulations of all test strains could not be killed at 24-hour, the tolerance assay was extended to 6-days to monitor the changes in the tolerance level of strains of different bacterial species under long term starvation. The experiment was designed such that, after the cells were subjected to 24-hour starvation, antibiotics at a concentration of ten times MIC (100 μg/ml ampicillin, 10 μg/ml gentamicin and 4 μg/ml ciprofloxacin) were added every second day to ensure that antibiotic concentration was maintained at a high level throughout the experiment. The size of the ampicillin tolerant bacterial sub-population remained stable at 10^8/ml throughout the 6-days treatment course. Effects of other β-lactam drugs such as meropenem were also tested, and the results were the same as that of ampicillin, suggesting that a subpopulation was tolerant to β-lactam drugs even after encountering starvation for a prolonged period. However, phenotypic tolerance to gentamicin and ciprofloxacin was found to vary significantly among strains of different bacterial species, indicating either that different degrees or nature of tolerance maintenance mechanisms are expressed in different bacterial species, or that protein synthesis and DNA replication functions remain active in some bacterial species.

**Figure 1.**
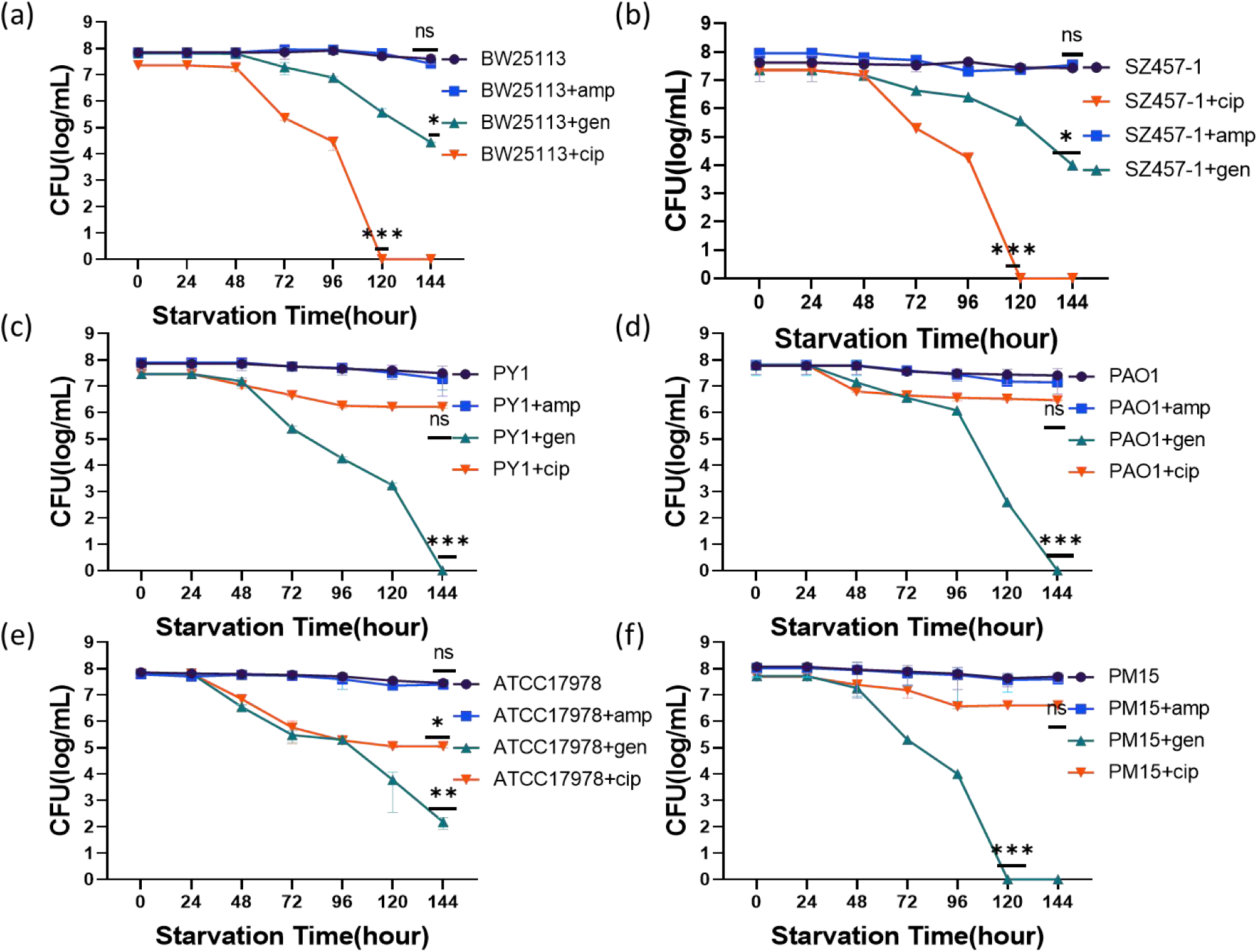
Tolerant subpopulations of different bacterial species can be killed by different antibiotics in the long term. amp, gen and cip stand for ampicillin, gentamicin and ciprofloxacin respectively. The tolerance level of *Escherichia coli* strain BW25113, SZ457-1, *Salmonella* strain PY1, *Pseudomonas aeruginosa* strain PAO1, *Acinetobacter baumannii* strain ATCC17978 and *Klebsiella pneumoniae* strain PM15 upon treatment with ampicillin, gentamicin and ciprofloxacin (10 times of MIC) is shown. Each value is presented as the mean, and error bar indicates SD. *, *P* < 0.05, **, *P* < 0.01, ***, *P* < 0.001, ns, *P* ≥ 0.05.

The tolerance level of *E. coli* strain BW25113 and SZ457-1 was found to decrease from 10^8 to 10^4 / ml upon treatment with gentamicin for six days, and that of *Pseudomonas aeruginosa* strain PAER00606, *Acinetobacter baumannii* strain ATCC17978 and CPC35 decreased to 10^3 / ml. Importantly, *Salmonella* strain PY1 and SR5, as well as the *Pseudomonas aeruginosa* strain PAO1 were completely killed by gentamicin in the 6-days experiment. On the other hand, *Klebsiella pneumoniae* strain PM15 and EH62 were found to be completely eradicated on Day 5 (Fig 1, S1). These findings indicate that bacterial strains that exhibit reduced antibiotic susceptibility do not necessarily exhibit higher tolerance levels. The data also further imply that protein synthesis remains active in tolerant subpopulations that form during starvation.

The degree of phenotypic tolerance to ciprofloxacin was also investigated and the results showed that the tolerance level of *Salmonella* strain PY1 and SR5, *Klebsiella pneumoniae* strain PM15 and *Pseudomonas aeruginosa* strain PAO1 only decreased from 10^8 to 10^7 upon treatment with ciprofloxacin for 6-days. The size of the tolerant subpopulation of the *Klebsiella pneumoniae* strain EH62, *Pseudomonas aeruginosa* strain PAER00606 *Acinetobacter baumannii* strain ATCC17978 and CPC35 was found to decrease to 10^5/ml. However, the tolerant subpopulations of *E. coli* strain BW25113 and SZ457-1 were completely eradicated on Day 6. Ciprofloxacin acts by inhibiting DNA replication in bacteria. Our finding confirm that DNA synthesis remains active in tolerant subpopulations of strains of specific bacterial species (Fig 1, S1). DNA replication is expected to be active only during bacterial cell proliferation, however, as bacteria do not replicate during starvation, we hypothesize that some types of uncharacterized DNA synthesis or repair activities occur in the tolerant subpopulations.

### Role of active DNA repair in mediating maintenance of antibiotic tolerance

As ciprofloxacin could eradicate *E. coli* BW25113 tolerant subpopulations completely after 6-days starvation, we further assessed the effects of ciprofloxacin on different types of tolerant subpopulations during the 6-days starvation episode and investigated the DNA synthesis activities in such subpopulations. The thymidine analog 5′-bromo-2′-deoxyuridine (BrdU) was used to label proliferating cells in a wide variety of species, including plant and mammalian cells, often in experiments designed to detect active DNA synthesis activity (Cecchini et al., 2012). First, we showed that almost every cell in the log phase control contained brdU, indicating that DNA replication was active. After encountering starvation for 24-hours, the fluorescence signals only slightly decreased, with that of the BW25113, SZ457-1, PM15 and ATCC17978 tolerant subpopulations decreasing from 60000 to 50000. Those of SR5 and PAER00606 were found to decrease from 40000 to 30000. At 72-hours starvation, the fluorescence intensity decreased sharply to around 20000, but was still detectable. At the end of 6-day starvation, the fluorescence intensity decreased to a level lower than 20000, and were relatively weak in tolerant subpopulations of all bacterial species (Fig S2a). This finding indicates that DNA replication in tolerant subpopulations was active at the beginning, and that this activity only declined gradually when starvation conditions persisted, with a low level of DNA synthesis activity still detectable upon starvation for 6-days. As the size of the tolerant subpopulations did not change during the 6-days starvation episode, and deletion of DNA replication genes did not affect the tolerance level, DNA replication could not have occurred in the tolerant subpopulations (Fig S3). To rule out the possibility that the fluorescence signals detectable in the antibiotic tolerant population was due to accumulation of the label molecules in the tolerant cells as a result of membrane damage that occurred during nutrient starvation, we extracted DNA from *E.coli* BW25113 at log phase, 24-hour, 72-hour and 144-hour starvation after staining, followed by washing out of the labelling molecules and measuring the fluorescence signals in the DNA samples. We were able to detect a high level of fluorescence signals in the DNA samples, the intensity of which only decreased gradually during the course of 144 hours starvation (Fig S3b). This observation indicates that the BrdU labels were incorporated into the newly synthesized DNA molecules, thereby confirming that DNA synthesis remained active in the antibiotic tolerant subpopulation.

We hypothesize that the DNA synthesis activities we observed in tolerant subpopulations involved repair of DNA strand damaged during nutrient starvation. To test this hypothesis, six genes which encode DNA repair functions were deleted individually to test whether DNA repair activity indeed occurs in tolerant subpopulations of *E. coli* BW25113 during starvation. As shown in Figure 2d, deletion of the *recB*, *recC*, and *uvrC* genes led to significantly larger extent of decrease in the population size of tolerant subpopulations than the wild type strain upon treatment with ciprofloxacin for 6-days. Consistently, results of the DNA synthesis labeling assay (Fig 2b) showed that the fluorescence signal of the *E.coli* BW25113::Δ*recB* and Δ*recC* knockout strains decreased significantly and almost disappeared at the end of the experiment. These findings further indicate that the DNA synthesis activity observed in the tolerant cells was attributed to DNA repair function, which remained active in tolerant subpopulations even after encountering starvation for 6-days. Apparently this function is essential in mediating onset and maintenance of antibiotic tolerance.

**Figure 2.**
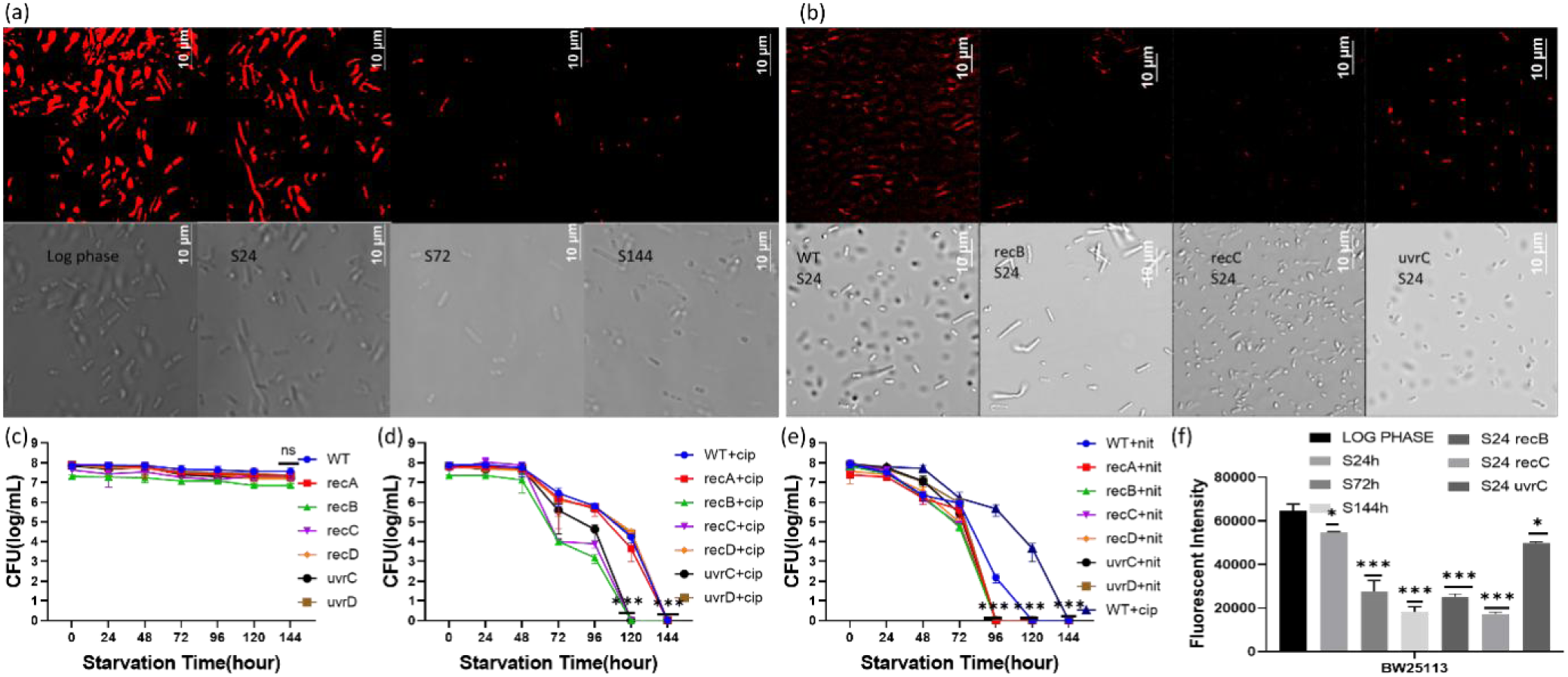
The role of DNA repair mechanisms in mediating expression and maintenance of antibiotic tolerance in *E. coli* strain BW25113. WT stands for *E. coli* BW25113 wild type strain; recA, recB, recC, recD, uvrC and uvrD stand for *E. coli* BW25113::Δ*recA*, Δ*recB*, Δ*recC*, Δ*recD*, Δ*uvrC* and Δ*uvrD* knockout strains, respectively; cip and nit stand for ciprofloxacin and nitrofurantoin respectively. (a) Results of DNA labeling assay of *E. coli* BW25113 in log phase, and tolerant subpopulations of such strains collected at 24, 72, 144-hour starvation. (b) Results of DNA labeling assy of the *E. coli* BW25113::Δ*recB*, Δ*recC* and Δ*uvrC* strains which had encountered nutrient starvation for 24-hours. (c) The viability of the six DNA repair gene knockout mutants upon encountering starvation for six days. (d) The level of tolerance of the six DNA repair gene knockout mutants to ciprofloxacin (10 X MIC) during a six-days treatment. (e) The level of tolerance of six DNA repair gene knockout mutants to nitrofurantoin (1X MIC) during a six-days treatment. (f) Fluorescence intensity of the six DNA gene knockout mutants recorded in DNA labeling assay upon encountering starvation for 24-hours. Each value is presented as the mean, and error bar indicates SD. *, *P* < 0.05, **, *P* < 0.01, ***, *P* < 0.001, ns, *P* ≥ 0.05.

Consistently, we found that the tolerant subpopulations were extremely susceptible to nitrofurantoin, a drug which is known to inhibit protein synthesis and induce DNA damages, thereby counteracting DNA repair activities in such subpopulations. We further tested the tolerance level of the mutants in which the *recA*, *recB*, *recC*, *recD*, *uvrC* and *uvrD* genes that encode DNA repair function, during nitrofurantoin treatment (Fig 2e), and found that the tolerant subpopulation of these gene knockout mutants were completely eradicated within 72 hours, even if the drug concentration was reduced to 1X MIC. This observation confirmed the importance of DNA repair activity in maintenance of starvation-induced antibiotic tolerance. Since tolerant subpopulations need to produce DNA repair enzymes to maintain DNA integrity, we also tested whether the combination of gentamicin and nitrofurantoin, and the combination of gentamicin and ciprofloxacin, acts synergistically to rapidly eradicate tolerant subpopulations. Our data showed that tolerant subpopulations of all bacterial species could be eradicated completely by the nitrofurantoin and gentamicin combination within 96 hours; in particular, tolerant subpopulations of *Acinetobacter baumannii* were totally killed within 72 hours. Likewise, gentamicin and ciprofloxacin were found to act synergistically on tolerant subpopulations by simultaneously inhibiting DNA repair functions and synthesis of enzymes responsible for such functions, with the killing effect of this drug combination being much stronger than that of gentamicin and ciprofloxacin alone (Fig 3).

**Figure 3.**
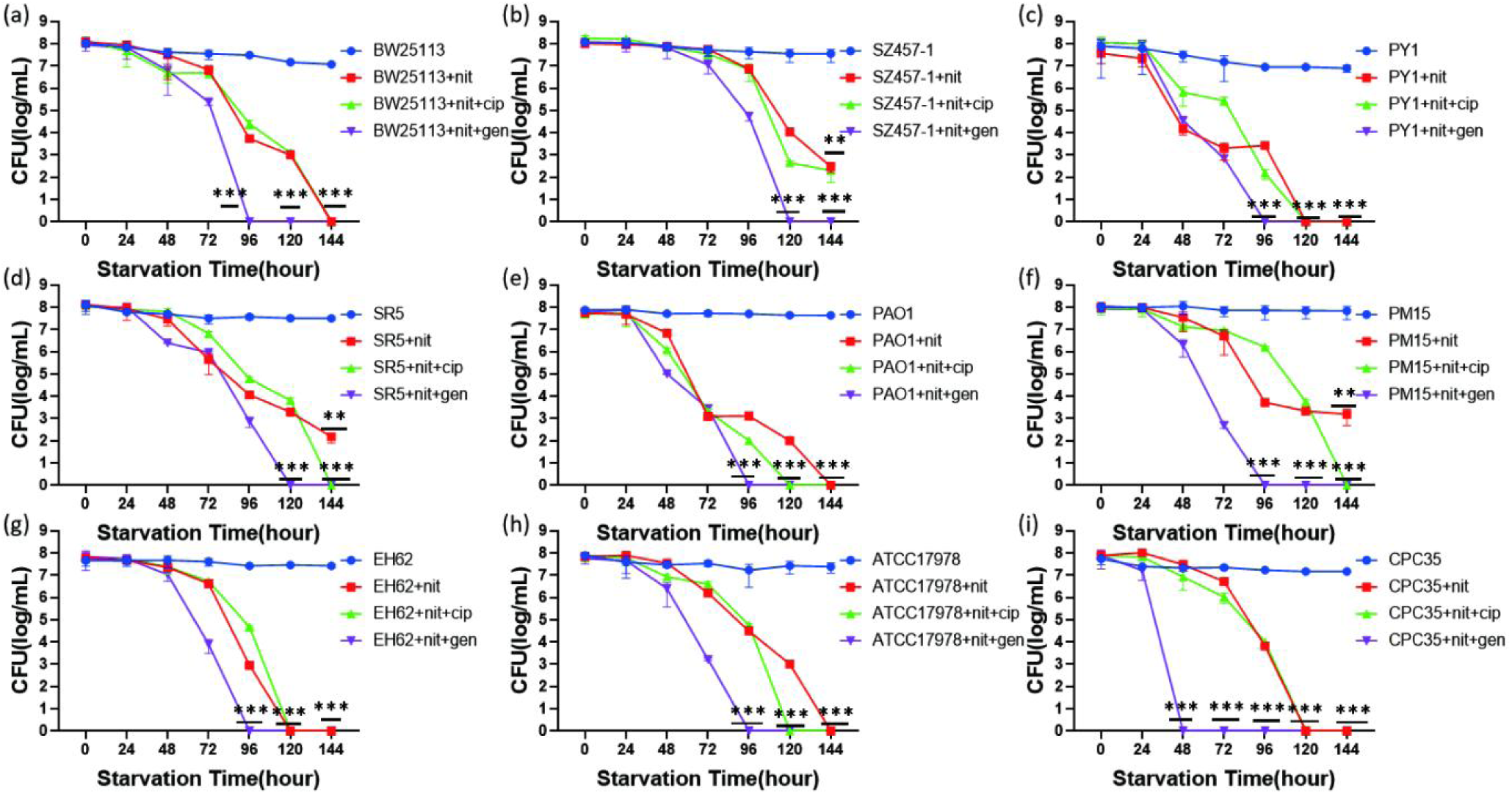
The combined anti-tolerance effects of nitrofurantoin and two antibiotics, gentamicin and ciprofloxacin. The abbreviations gen, cip and nit stand for gentamicin, ciprofloxacin and nitrofurantoin respectively. (a) The level of tolerance of *Escherichia coli* BW25113 to nitrofurantoin (1X MIC), nitrofurantoin (1X MIC) and ciprofloxacin (10X MIC), nitrofurantoin (1X MIC) and gentamicin (10X MIC) during a 144-hours treatment. (b) The level of tolerance of *Escherichia coli* SZ457-1 to nitrofurantoin (1X MIC), nitrofurantoin (1X MIC) and ciprofloxacin (10X MIC), nitrofurantoin (1X MIC) and gentamicin (10X MIC) during a 144-hours treatment. (c) The level of tolerance of *Salmonella* PY1 to nitrofurantoin (1X MIC), nitrofurantoin (1X MIC) and ciprofloxacin (10X MIC), nitrofurantoin (1X MIC) and gentamicin (10X MIC) during a 144-hours treatment. (d) The level of tolerance of *Salmonella* SR5 to nitrofurantoin (1X MIC), nitrofurantoin (1X MIC) and ciprofloxacin (10X MIC), nitrofurantoin (1X MIC) and gentamicin (10X MIC) during a 144-hours treatment. (e) The level of tolerance of *Pseudomonas aeruginosa* PAO1 to nitrofurantoin (1X MIC), nitrofurantoin (1X MIC) and ciprofloxacin (10X MIC), nitrofurantoin (1X MIC) and gentamicin (10X MIC) during a 144-hours treatment. (f) The level of tolerance of *Klebsiella pneumoniae* PM15 to nitrofurantoin (1X MIC), nitrofurantoin (1X MIC) and ciprofloxacin (10X MIC), nitrofurantoin (1X MIC) and gentamicin (10X MIC) during a 144-hours treatment. (g) The level of tolerance of *Klebsiella pneumoniae* EH62 to nitrofurantoin (1X MIC), nitrofurantoin (1X MIC) and ciprofloxacin (10X MIC), nitrofurantoin (1X MIC) and gentamicin (10X MIC) during a 144-hours treatment. (h) The level of tolerance of *Acinetobacter baumannii* ATCC17978 to nitrofurantoin (1X MIC), nitrofurantoin (1X MIC) and ciprofloxacin (10X MIC), nitrofurantoin (1X MIC) and gentamicin (10X MIC) during a 144-hours treatment. (i)The level of tolerance of *Acinetobacter baumannii* CPC35 to nitrofurantoin (1X MIC), nitrofurantoin (1X MIC) and ciprofloxacin (10X MIC), nitrofurantoin (1X MIC) and gentamicin (10X MIC) during a 144-hours treatment. Each value is presented as the mean, and error bar indicates SD. *, *P* < 0.05, **, *P* < 0.01, ***, *P* < 0.001, ns, *P* ≥ 0.05.

### Protein synthesis functions are also required for maintenance of antibiotic tolerance

As gentamicin was found to exhibit a certain degree of killing effects on tolerant subpopulations of all the 11 Gram-negative bacterial strains, we investigated whether the tolerant subpopulations of these species continued to synthesize protein during starvation. L-Glutamic acid γ-(7-amido-4-methylcoumarin), in which the amino acid glutamic acid is linked to a fluorescent dye, was used to label newly synthesized protein strands in the tolerant subpopulations. Figure 4a and S4 showed that the fluorescence signals detectable in the log phase cells were strong, and that almost every cell contained of L-Glutamic acid γ-(7-amido-4-methylcoumarin), indicating that protein synthesis was active. After encountering starvation for 24-hours, the fluorescence signals decreased significantly, with the fluorescence intensity of the BW25113 and PM15 tolerant subpopulations decreasing from 200000 to 100000, while those of SZ457-1 and ATCC17978 decreased from 200000 to 150000. On the other hand, the fluorescence signals of tolerant subpopulations of the strain SR5 and PAER00606 were found to decrease from 250000 to 100000. At 72-hour starvation, the fluorescence intensity of all samples generally decreased to around 50000, but was still detectable. At the end of the 6-days starvation period, the fluorescence intensity decreased to a level below 50000, and was relatively weak in all bacterial species. To confirm that the fluorescence signals observed in the tolerant cells were not due to accumulation of the label molecules in such cells as a result of possible membrane damage under the test condition, we extracted protein of *E.coli* strain BW25113 at log phase, 24-hour, 72-hour and 144-hour starvation after staining, followed by washing out of the labeling molecules and measuring the fluorescence signals of the protein samples. The results showed that fluorescence signals were detectable in the protein samples, and that the signal level only decreased gradually during starvation (Fig S4b). This finding indicates that protein synthesis in tolerant subpopulations remained active and that the labeling molecules were incorporated into newly synthesized proteins. Nevertheless, the protein synthesis activity decreased gradually when starvation conditions persisted, yet a low level of protein synthesis activity was still detectable in tolerant cells which had encountered starvation for 6-days. In order to investigate the types of protein synthesized in tolerant subpopulations during starvation, comparative RNA sequencing analysis of tolerant subpopulations which had encountered starvation for six days and bacteria at log phase were performed. The P value of GO pathways (Fig 4b) showed that the expression status of several metabolic pathways differed significantly between the tolerant subpopulations and log phase cells. Among them were genes that encode the structural components of ribosome, protein-bearing complex, membrane protein complex and respiratory electron transport chain components (Figure S5a, 5b, 5c and 5d). The 22 most up-regulated genes in the tolerant subpopulations were studied by performing gene deletion experiments. Deletion of such genes, namely the *kdpB*, *rpoS*, *yeaG*, *aceA*, *kgtP*, *clpA* and *clpX* genes, was found to cause significant shrinkage of the tolerant population upon treatment with gentamicin for 6 days, when compared to the wild type control (Figure 4c). Consistently, the fluorescence signals decreased significantly in *E.coli* BW25113::Δ*kdpB*, Δ*rpoS,* Δ*yeaG*, Δ*kgtP* and Δ*clpA* knockout strains at 24h starvation, indicating that products of these genes were responsible for synthesis of proteins required for maintenance of antibiotic tolerance in the tolerant subpopulations that form during starvation.

**Figure 4.**
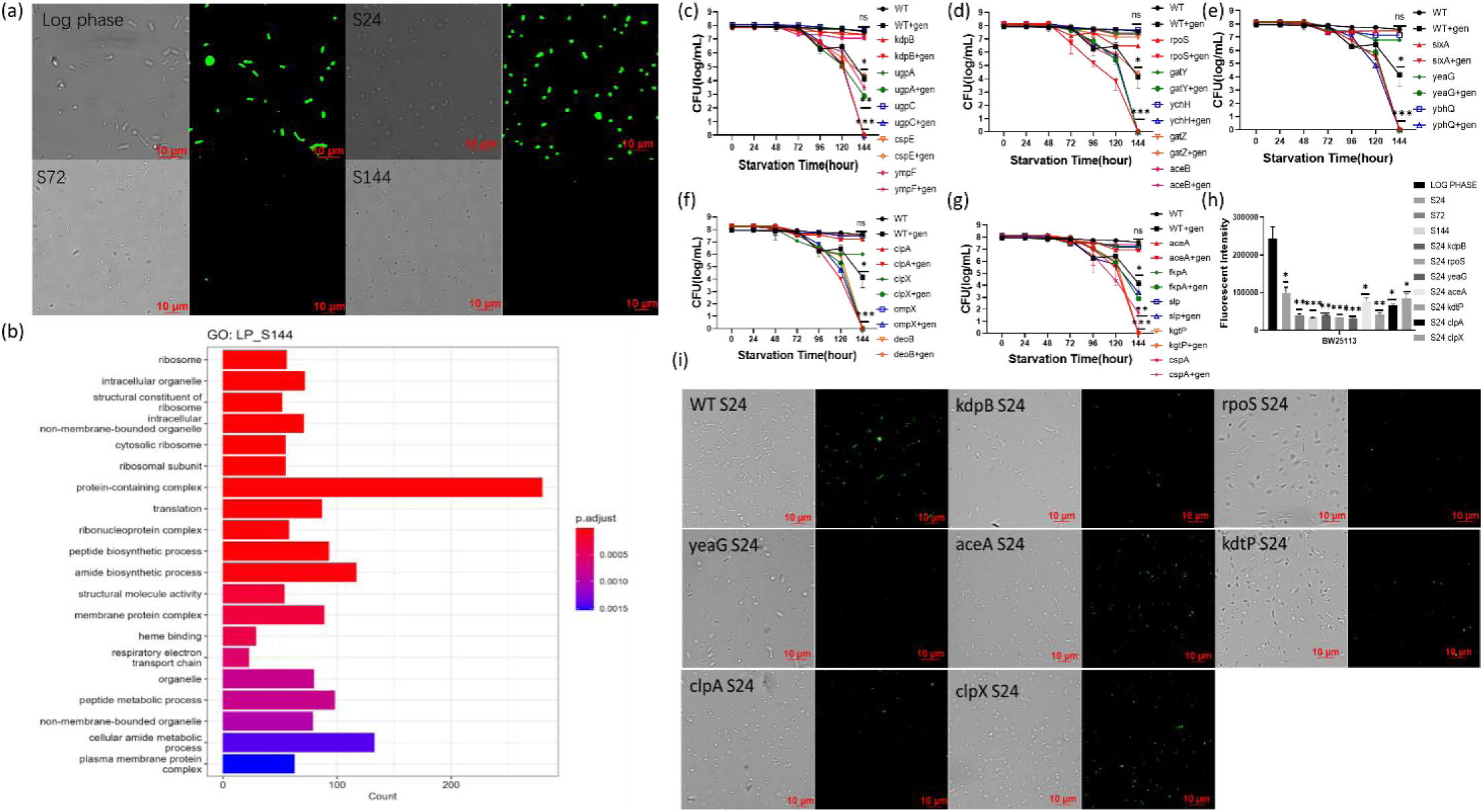
The role of protein synthesis mechanism in mediating expression and maintenance of antibiotic tolerance in *E. coli* strain BW25113. WT stands for *E. coli* BW25113 wild type strain; kdpB, rpoS, yeaG, aceA, kdtP, clpA and clpX stand for *E. coli* BW25113::Δ*kdpB*, Δ*rpoS*, Δ*yeaG*, Δ*aceA*, Δ*kdtP*, Δ*clpA* and Δ*clpX* gene knockout strains; gen stands for gentamicin. (a) Protein labeling assay results of *E. coli* BW25113 strain at log phase, and tolerant subpopulations of these strains which had encountered starvation for 24, 72, 144-hours. (b) The differences between log phase bacteria and tolerant subpopulations collected at 144 hours-starvation in terms of GO pathways. (c, d, e, f, g) The gentamicin tolerance level of strains in which genes involved in protein synthesis were deleted. (h) Fluorescence intensity of protein synthesis gene knockout strains in protein labeling assay upon 24-hour starvation. (i) Images of the gene knockout strains taken during protein labeling assay. Each value is presented as the mean, and error bar indicates SD. *, *P* < 0.05, **, *P* < 0.01, ***, *P* < 0.001, ns, *P* ≥ 0.05, WT, wild type strain.

### Changes in other physiological parameters of tolerant subpopulations upon prolonged starvation

We next performed NADH, ROS, efflux, ATP and membrane potential assays in tolerant subpopulations which had encountered starvation for 24, 72 and 144h, with log phase cells as control. The results helped explain why tolerant subpopulations were increasingly susceptible to antibiotics during the course of a 6-days treatment even though DNA, protein and cell wall synthesis activities decreased during this period. First, ATP level was found to decrease sharply at 24h (Figure S6), impeding cell wall and DNA synthesis. Hence bacteria became dormant and unresponsive to antibiotics at 24h. Second, the NADH level was found to decrease gradually during the course of starvation (Figure 5f, S7). This phenomenon is associated with a decline in oxidative phosphorylation activities and hence a lower level of PMF and efflux activities. Third, a lower energy level would also affect efflux. These events in turn cause Nile red and antibiotics to accumulate inside the bacterial cells (Figure 5d, S8), thereby exerting their inhibitory effects on tolerant subpopulations which still exhibited a low level of DNA, protein and cell wall synthesis activities. Besides, ROS were also found to accumulate inside the tolerant cells, presumably due to decrease in expression of enzymes such as superoxide dismutase, which neutralize ROS, thereby resulting in oxidative damages of cellular components of the tolerant cells (Figure S9). In order to further study the functional role of ATP generation machinery of bacteria in maintenance of the tolerance phenotypes, the effect of deletion of eleven genes involved in ATP synthesis in *E. coli* strain BW25113 was tested. Upon ciprofloxacin treatment (Figure S10), the tolerance phenotypes of these gene knockout strains were comparable with the wild type, indicating that a lack of ATP synthesis function does not severely affect DNA repair activity in the tolerant subpopulations. Upon treatment with gentamicin (Figure 5, S11), however, the size of the tolerant population of the *E.coli* BW25113::Δ*atpE*, Δ*atpH,* and Δ*atpI* knockout strains decreased significantly, suggesting that ATP is required for protein synthesis in the tolerant subpopulations. These three knockout strains were then tested in protein labeling assay upon encountering starvation for 24h. The fluorescence signals of the *atpH* and *atpI* mutants were found to decrease significantly when compared to the wild type strain, confirming that these three genes contributed to protein synthesis and hence stress defense in the tolerant subpopulations that form during nutrient starvation.

**Figure 5.**
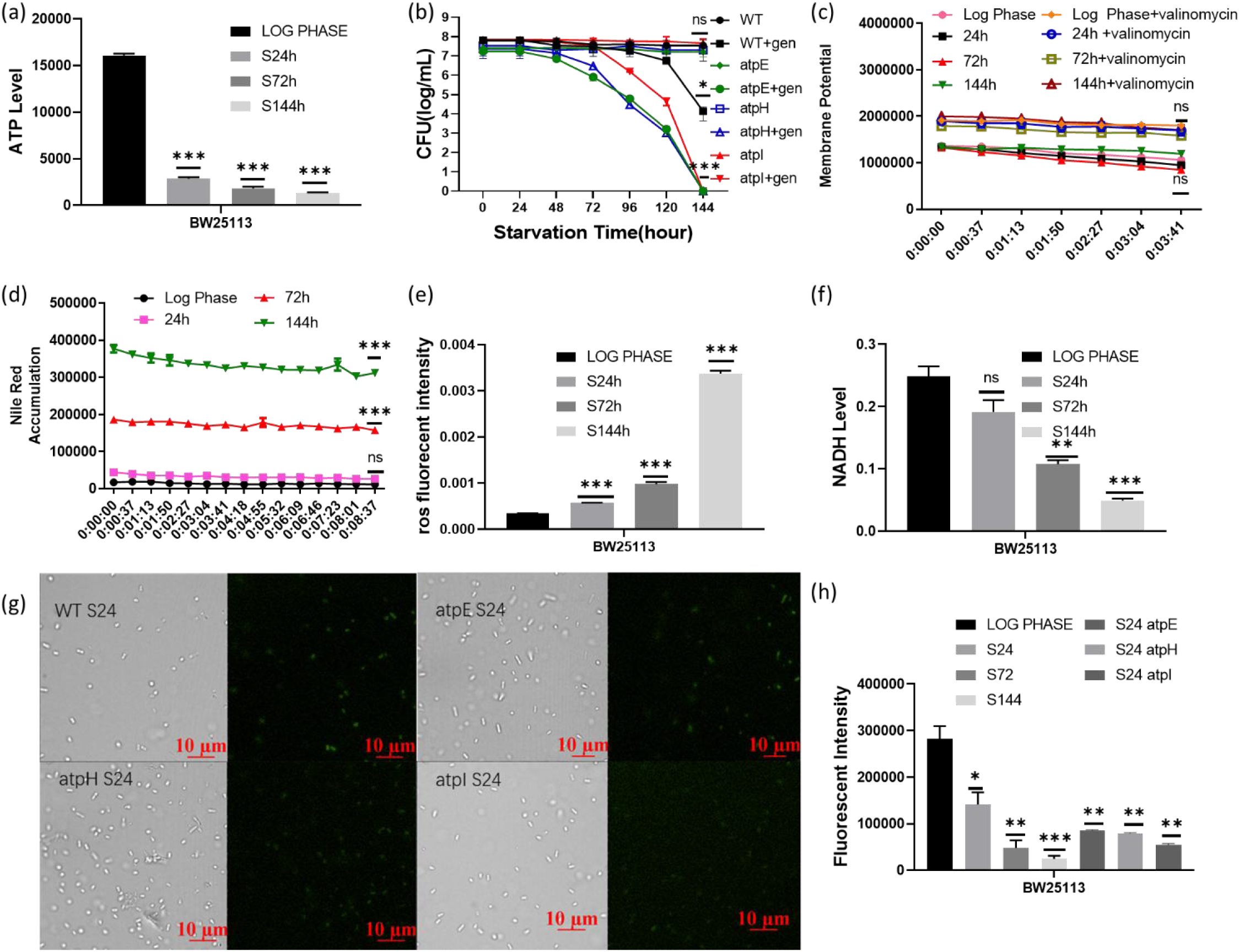
Other key cellular mechanisms that mediate expression and maintenance of antibiotic tolerance in *E. coli* BW25113. WT stands for *E. coli* BW25113 wild type strain; atpE, atpH, and atpI stand for the *E.coli* BW25113::Δ*atpE*, Δ*atpH* and Δ*atpI* gene knockout strains; gen stands for gentamicin. (a) ATP level of *E. coli* BW25113 at log phase, and that of tolerant subpopulations collected during 24, 72, 144-hours starvation. (b) Changes in gentamicin tolerance level of *E. coli* BW25113::Δ*atpE*, Δ*atpH* and Δ*atpI* knockout strains during 144 hours treatment. Membrane potential (c), Nile red accumulation (d), ros level (e) and NADH level (f) in *E. coli* strain BW25113 at log phase, and the corresponding tolerant subpopulations collected at 24, 72, 144-hours starvation. (g) Images of *E. coli* BW25113::Δ*atpE*, Δ*atpH* and Δ*atpI* gene knockout strains taken during protein labeling experiments. (h) Fluorescence intensity of ATP synthesis gene knockout strains recorded in protein labeling assay upon encountering 24-hours starvation. Each value is presented as the mean, and error bar indicates SD. *, *P* < 0.05, **, *P* < 0.01, ***, *P* < 0.001, ns, *P* ≥ 0.05, WT, wild type strain.

## Discussion and Summary

In this work, we showed that tolerant subpopulations of different bacterial species exhibited varied abilities to maintain their tolerance phenotypes upon antimicrobial treatment. For example, treatment with gentamicin for six days only led to reduction in the size of tolerant subpopulations of *E. coli* strain BW25113, SZ457-1, and *Pseudomonas aeruginosa* strain PAER00606 from 10^8 to 10^4 / ml, but was able to completely eradicate tolerant subpopulations of the *Salmonella* strain PY1 and SR5, the *Pseudomonas aeruginosa* strain PAO1, as well as the *Klebsiella pneumoniae* strain EH62. The degree of tolerance of various bacterial species to ciprofloxacin treatment was also different. The population size of tolerant subpopulations of *Salmonella* strain PY1 and SR5, *Klebsiella pneumoniae* strain PM15 and *Pseudomonas aeruginosa* strains PAO1 only decreased from 10^8 to 10^7 upon treatment with ciprofloxacin for six days, however, those of *E. coli* strain BW25113 were completely eradicated. The significant discrepancy in the level of tolerance of each bacterial species to gentamicin and ciprofloxacin is intriguing. Nevertheless, this phenomenon indicates that each bacterial species expresses unique yet active and complex underlying physiological functions and defensive mechanisms that define their antibiotic tolerance phenotype. On the other hand, our data also indicate that the tolerance level of the test bacterial strains did not correlate with the degree of susceptibility to various antibiotics. In other words, drug resistant strains may be eradicated by antibiotics when their physiological status is switched to the tolerance mode. This is due to the fact that the resistance mechanisms of the drug resistant strains do not contribute to maintenance of stress or antibiotic tolerance. This finding has important implication as it provides the opportunity of targeting drug resistant organisms by suppressing their stress tolerance responses.

It is conceivable that antibiotic tolerant subpopulations of various bacterial species are susceptible to protein synthesis inhibitors such as aminoglycosides in the long term as our recent studies showed that they need to synthesize specific proteins to maintain PMF and efflux activities, as well as other essential physiological functions, in order to survive against adverse environmental conditions (Wan et al., 2023; Wan et al., 2021). Yet it is surprising to find that some bacterial species only exhibited very low level of tolerance to ciprofloxacin, which is one of the fluoroquinolone antibiotics which can inhibit DNA replication and cause DNA breakage in bacteria. Such finding suggests that DNA synthesis is still active in the tolerant subpopulations. Our labeling studies indeed showed that DNA synthesis remained detectable in bacteria which had encountered starvation for six days. However, bacteria do not multiply during starvation and are therefore not expected to undergo DNA replication. The DNA synthesis activities could only be due to the need to repair damaged DNA in the tolerant subpopulations. A study by Völzing et al. showed that recovery of stationary phase tolerant subpopulations required DNA repair functions (Volzing & Brynildsen, 2015). Our study confirmed the role of DNA repair in mediating expression of antibiotic tolerance by testing mutants in which DNA repair genes were deleted, as such mutants exhibited significantly lower ability to maintain phenotypic tolerance to ciprofloxacin. We also confirmed that the tolerant cells of both the wild type strain and DNA repair gene deletion mutants were effectively eradicated by ciprofloxacin by performing live /dead staining assay (Fig S12). Consistently, the drug nitrofurantoin was found to be extremely effective in eradicating bacterial tolerant cells. In particular, this drug acts synergistically with gentamicin to kill tolerant subpopulations in a highly efficient manner. As nitrofurantoin is postulated to inflict DNA damage or inhibit DNA repair and protein synthesis, this finding confirms that DNA repair functions are essential for prolonged survival of bacterial antibiotic tolerant subpopulations. Specific DNA repair mechanisms in the tolerant subpopulations and the degree by which such mechanisms confer fluoroquinolone tolerance need to be further investigated.

Regardless of the nature of mechanisms involved in DNA repair, our findings are consistent with results of RNA sequencing, protein labeling assay and tolerance assays with gene deletion mutants, which consistently showed that tolerant subpopulations synthesize a range of proteins that play a role in maintaining the tolerance phenotype. This work adds DNA repair proteins to the list of proteins required for maintaining phenotypic antibiotic tolerance in bacteria. It should also be noted that some of genes involved in protein synthesis in tolerant subpopulations are expressed at a level even higher than that of the log phase cells. These genes include *kdpB*, *rpoS*, *yeaG*, *kgtP* and *clpA*. It is necessary to investigate whether these genes mainly play a role in mediating protein synthesis in tolerant subpopulations but not exponentially growing bacterial cells. Apart from the unique sets of genes which appear to encode DNA repair and protein synthesis functions in tolerant subpopulations but not exponentially growing cells, we suspect that tolerant subpopulations also produce other proteins or enzymes that play a specific role in supporting continuous expression of the tolerance phenotypes. Another possible area where unique protein products need to be synthesized in tolerant subpopulations is the bacterial cell wall. Our work showed that β-lactam drugs could not kill tolerant subpopulations despite the fact that cell wall synthesis activities were detectable. It remains to be seen whether specific proteins to which known β-lactam cannot bind tightly are involved in cell wall synthesis in tolerant subpopulations.

One interesting observation in this study is that, despite detection of a relatively high DNA repair and protein synthesis activity when tolerant subpopulations first formed when they encountered nutrient starvation, tolerance to ciprofloxacin and gentamicin was at the highest level at that early stage; yet the strength of tolerance to these two drugs gradually decreased when starvation conditions persisted and the DNA and protein synthesis activities became lower. The phenomenon can be explained by the finding that ATP level and efflux activities also declined gradually during the course of starvation. ATP was found to be required for synthesis of proteins required for maintaining the tolerance phenotypes in tolerant subpopulations, as well as for DNA repair; hence the level of ATP, protein synthesis and DNA repair activities decreased simultaneously when starvation conditions became long-lasting. Losing the ability to synthesize proteins by itself has the effect of weakening various tolerance maintenance mechanisms, including the essential DNA repair functions, resulting in gradual reduction tolerance level. Our previous study showed that tolerant subpopulations need to maintain proton motive force (PMF) and synthesize a number of efflux pumps, presumably to expel toxic metabolites and antibiotics during nutrient starvation (Wan et al., 2023; Wan et al., 2021). Both processes also require synthesizing specific proteins such as PspA. Failure to maintain PMF will also affect efflux. Lower PMF level and reduced efflux activity would lead to antibiotic accumulation in tolerant cells. As tolerant subpopulations maintain a range of physiological functions even after six days of starvation, gradual accumulation of antibiotics would exert an increasingly strong killing effect on these bacterial cells. Although DNA repair and protein synthesis activities gradually wind down as nutrient starvation condition persist, these activities are subjected to inhibition by the antibiotics that start to accumulate intracellularly. Once this happens, these events would enter a vicious cycle in which all essential tolerance maintenance functions would stop and the tolerant subpopulations would die. Finally, we also found that tolerant subpopulations need to neutralize reactive oxygen species (ROS) in order to survive. Hence, prolonged starvation or treatment with antibiotics that inhibit protein synthesis would also affect expression of enzymes such as superoxide dismutase that play a role in ROS neutralization, thereby resulting in a reduced tolerance level in tolerant subpopulations.

To conclude, our works show that bacterial antibiotic tolerant subpopulations that form during nutrient starvation are not physiologically dormant, instead a wide range of cellular activities associated with various repair and defense functions remained detectable after six days of starvation, even though such activities gradually subside during the process. These activities, which involve antibiotic-sensitive DNA repair and protein synthesis activities, render tolerant subpopulations increasingly susceptible to antibiotics as weakening energy production and efflux mechanisms lead to accumulation of antibiotics in the intracellular compartment and cause further inhibition of the drug-sensitive DNA repair and protein synthesis activities. These findings of drug targets indicate that various commonly used antibiotics can be incorporated in design of new strategies to eradicate antibiotic tolerant subpopulations to prevent occurrence of chronic and recurrent infections.

## Methodology

Detailed methods are provided in the online version of this paper and include the following:

### KEY RESOURCES

Luria-Bertani (LB) broth was used for all cultures unless stated otherwise. Standard LB agar (Difco, Leeuwarden, The Netherlands) was used in testing the proportion of the bacterial population that survived in the antibiotic persistence assay. MH broth were purchased from Qingdao Hope Bio-Technology Co., Ltd. Ampicillin, gentamicin, ciprofloxacin and nitrofurantoin were purchased from Sigma (St. Louis, MO). DiSC3(5) was purchased from Invitrogen. BrdU and HADA were obtained from ThermoFisher and Sigma, respectively. NaCl, KCl, and MgCl_2_ was purchased from UNI-CHEM.

### EXPERIMENTAL MODEL AND SUBJECT DETAILS

#### Bacterial strains

| Strains | Source |
| --- | --- |
| *Escherichia coli* BW25113 | Coli Genetics Stock Center |
| *Escherichia coli* SZ457-1 | Shen Zhen Food Industry |
| *Klebsiella pneumoniae* PM15 | Hong Kong Hospital |
| *Klebsiella pneumoniae* EH62 | Hong Kong Hospital |
| *Salmonella typhimurium* PY1 | Hong Kong Hospital |
| *Salmonella typhimurium* SR5 | Hong Kong Hospital |
| *Pseudomonas aeruginosa* PA01 | DSMZ |
| *Pseudomonas aeruginosa* PAER00606 | Hong Kong Hospital |
| *Acinetobacter baumannii* ATCC17978 | ATCC |
| *Acinetobacter baumannii* CPC35 | Hong Kong Hospital |
| Gene knockout strains | Coli Genetics Stock Center |

### METHODS

#### Determination of Minimal Inhibitory Concentrations (MICs)

The MIC results of ampicillin, gentamicin and ciprofloxacin against *Escherichia coli* BW25113 (Grenier et al., 2014), SZ457-1, *Salmonella* PY1 (Chen et al., 2007), SR5 (Moosdeen & Cheong, 1989), *Klebsiella pneumoniae* PM15, EH62, *Acinetobacter baumannii* ATCC17978, CPC35 (Mayer et al., 2018) and *Pseudomonas aeruginosa* PAER00606, PAO1 (Stover et al., 2000) were determined by using the Mueller Hinton Broth (MHB) (BD Difco, America). Mueller Hinton agar plates containing various concentrations of ampicillin, gentamicin and ciprofloxacin were inoculated with the test strains and grown at 37°C for 16-22 hours. The results were compared with that of the standard strain *Escherichia coli* ATCC25922 according to CLSI guideline (Wayne, 2011).

#### Starvation-induced tolerance assay

Bacterial populations containing *Escherichia coli* BW25113, SZ457-1, *Salmonella* PY1, SR5, *Klebsiella pneumoniae* PM15, EH62, *Acinetobacter baumannii* ATCC17978, CPC35 and *Pseudomonas aeruginosa* PAER00606, PAO1 were subjected to starvation stress followed by antibiotic tolerance assay. Briefly, bacterial culture at mid-log phase was subjected to centrifugation (6000g, 5min), followed by removal of supernatant and resuspension of the pellet in 0.85% saline. The test antibiotic (ten times of MIC) was added to the cell suspension and then every second day for up to 6 days. The cell suspension was then incubated at 37℃ and shaken at 250 rpm. Survival rate of the bacteria in the cell suspension was recorded by recording the CFU (colony-forming unit) daily. Every assay was repeated three times. Live/dead cell imaging kit (ThermoFisher) was used to confirm that the tolerant cells of test strains were eradicated by the test antibiotic.

#### DNA labeling in gene knockout strains

In order to assess DNA synthesis activity in tolerant subpopulations under 6-day starvation, *E. coli* BW25113, SZ457-1, *Klebsiella pneumoniae* PM15, *Salmonella* SR5, *Pseudomonas aeruginosa* PAER00606 and *Acinetobacter baumannii* ATCC17978 were grown to log phase (OD= 0.2), 0.85% saline was used to wash the cells and remove LB broth to create starvation condition as described in the tolerance assay. Starvation condition was kept for 6 days to allow formation of tolerant subpopulations, and tolerant subpopulations collected at three time points, 24h, 72h and 144h were subjected to analysis by fluorescence microscopy, with log phase bacteria included a control. At each time point, the tolerant cells were centrifuged at 6500 g for 5 min, after which supernatant was discarded, and a solution containing 10 μM brdU was used to resuspend the bacteria, which were then stained at room temperature for 2 hours and kept in dark. PBS buffer was then used to wash the cells and remove the fluorescence dye to prevent the background from being too bright so that the bacterial cells cannot be observed. The wavelength of brdU is 523nm and a EMCCD camera was used to record the fluorescence signal. The images were analyzed by the ImageJ software (Fiji). To measure the fluorescence intensity, samples were added into a 96-well black cell culture plate and a plate reader was used to record the fluorescence signals. In order to that the brdU molecules were incorporated into the newly synthesized DNA strand rather than being accumulated in the bacterial cell due to membrane damage, DNA was extracted from *E.coli* strain BW25113 at different time points during the tolerance assay by using a DNA extraction kit (ThermoFisher) after the staining step, followed by washing of the cells to remove the brdU labels. The fluorescence signal of each sample was measured by a plate reader.

#### Protein labeling in gene knockout strains

In order to observe protein synthesis activity in tolerant subpopulations under 6-day starvation, *E. coli* BW25113, SZ457-1, *Klebsiella pneumoniae* PM15, *Salmonella* SR5, *Pseudomonas aeruginosa* PAER00606 and *Acinetobacter baumannii* ATCC17978 were grown to log phase (OD= 0.2), and washed with 0.85% saline to create starvation condition, which was maintained for 6 days to allow formation of tolerant subpopulations; the tolerant subpopulations were collected at three time points, 24h, 72h and 144h, and subjected to analysis by fluorescence microscopy, with log phase bacteria as control. Briefly, cells collected at each time point were centrifuged at 6500 g for 5 min, after which the supernatant was discarded, and 100µM fluorescent glutamic acid was added. The bacteria were then incubated for 30mins and kept in dark. PBS buffer was then used to wash the cells and remove the fluorescent dye. The wavelength of fluorescent glutamic acid is 470nm and a EMCCD camera was used to record the fluorescence signal. The images were analyzed by the ImageJ software (Fiji). To measure the fluorescence intensity, the samples were added into a 96-well black cell culture plate and a plate reader was used to record the fluorescence signals. In order to confirm that the glutamic acid label was incorporated into newly synthesized protein molecules rather than being accumulated in the bacterial cell as a result of membrane damage, protein was extracted from *E.coli* strain BW25113 at different time points as mentioned above by using the B-PER protein extraction reagent which contained lysozyme, PIC (protease inhibitor) and DNase after the staining step, followed by removal of the excess labels and measurement of the fluorescence signal of each sample by using a plate reader.

#### ATP assay

Overnight culture was prepared; 50 μl overnight culture were then added into 5 ml LB broth until a cell density of OD value of 0.2 was reached; the diluted culture was then incubated at 37℃, followed by washing and resuspending in 0.85% saline; the bacterial suspension was then incubated for up to 144h to create starvation stress; bacteria at log phase were used as control. Bacteria collected at different time points were then lysed for determination of the ATP level. At the same time, 1 ml 1× Reaction Buffer was prepared by adding 50 ml of 20× Reaction Buffer (Component E) to 950 ul of deionized water; 1 ml of 10mM D-lucferin stock solution was obtained by adding the buffer to one vial of D-lucferin (component A). The solution was protected from light until use. 1.62 ml of dH_2_O was added to a bottle containing 25 mg of DTT (component C). The solution was then divided into 10 160 μl volumes and kept on ice until use. In the next step, 5 mM ATP solution was prepared as ATP standard (component D). The reaction solution was as follows, 8.9 ml dH2O, 0.5 ml 20× Reaction Buffer (Component E), 0.1 ml 10mM D-lucferin and 2.5 μl of firefly luciferase 5 mg/ml stock solution. The reaction solution was mixed gently, and then added to a ATP standard to plot a standard curve, reaction solution of the same volume was added into samples and a plate reader was used to record the results.

#### NADH assay

50 μl overnight culture of each test strain were then added into 5 ml LB broth, and incubated at 37℃ until OD of 0.2 was reached; 0.85% saline was then used to wash and suspend the bacteria, followed by incubation for 144h to create starvation stress; bacteria at log phase were included as control. In the next step, bacteria collected at different time points were concentrated to OD of 0.4, followed by addition of 100 μl NADH extraction buffer, heat extraction at 60℃ for 5 min, and addition of 20 μl Assay Buffer and 100 μl NAD extraction buffer to neutralize the extracts. The solution was then vortexed and centrifuged at 14,000 rpm for 5 min, and the supernatant was collected for the assay. Standard sample of 0, 3, 6 and 10 μl NAD was prepared by mixing distilled water and Premix. The reagent was prepared by mixing 50 μl Assay Buffer, 1 μl Enzyme A, 1 μl Enzyme B, 14 μl Lactate and 14 μl MTT. 80 μl working reagent and 40 μl samples were added to each well; OD of each sample was recorded by a plate reader.

#### Efflux activity assay

50 μl overnight culture was added into 5 ml LB broth, and incubated at 37℃ until OD of 0.2 was reached; 0.85% saline was used to wash and suspend the bacteria, followed by incubation for 144h to create starvation stress; bacteria at log phase was included as control. Bacteria collected at different time points were then concentrated to OD of 0.4 and stained with 5μM of Nile red for 3 hours at 37℃. After staining, thebacteria were washed and resuspended with PBS and 1mM MgCl_2_, and then transferred to a 96-well black plate. The fluorescence intensity in each well was measured by a plate reader at 560nm and 655nm.

#### Membrane potential assay

50 μl overnight culture was added into 5 ml LB broth and incubated at 37℃ until OD 0.2 was reached; 0.85% saline was used to wash and suspend the bacterial cells, followed by incubation for 144h to create starvation stress; bacteria at log phase were included as control. DiSC3(5) was used as the membrane potential-sensitive probe, KCl was used to create proton and valinomycin which can destroy PMF was used as a positive control. KCl and DiSC3(5) were added until a final concentration of 100mM and 1μM was respectively reached, followed by incubation at room temperature for 15 mins in dark to allow the dye to penetrate through the outer membrane and produce a quenching effect. Valinomycin (1μM) was then added to the positive control group. The fluorescence reading was monitored by using a Clariostar Microplate Reader (BMG LABTECH) at an excitation wavelength of 622+10 nm and an emission wavelength of 670+10 nm for 10mins. Upon depolarization, the dye was rapidly released into the medium, resulting in dequenching and facilitating detection fluorometrically (Te Winkel et al., 2016)

#### ROS assay

50 μl overnight culture was added into 5 ml LB broth and then incubated at 37℃ until a cell density of OD 0.2 was reached; 0.85% saline was used to wash and suspend the bacteria, followed by incubation for up to 144h to create starvation stress; bacteria at log phase were included as control. Bacteria collected at different time points were incubated with DCFDA (Invitrogen) for 30mins. After staining, the bacteria were washed and resuspended by PBS and transferred to a 96-well black plate. The fluorescence intensity of each sample was measured by a plate reader at 485nm and 535nm.

### QUANTIFICATION AND STATISTIC ALANALYSIS

Details of all statistical analyses performed in this study are described in the respective figure legends.

## Acknowledgments

The work was supported by the Guangdong Major Project of Basic and Applied Basic Research (2020B0301030005) and the grant from Research Grant Council of Hong Kong Government (T11-104/22-R), as well as the grant from Health and Medical Research Fund (20190802).

---

| MIC | Amp (µg/ml) | Gen (µg/ml) | Cip (µg/ml) | Nitrofurantoin (µg/ml) |
| --- | --- | --- | --- | --- |
| *Escherichia coli* BW25113 | 8 | 1 | 0.125 | 32 |
| *Escherichia coli* SZ457-1 | 8192 | 1 | 0.125 | 256 |
| *Klebsiella pneumoniae* PM15 | 1024 | 1 | 0.015 | 64 |
| *Salmonella typhimurium* PY1 | 8 | 1 | 0.015 | 256 |
| *Salmonella typhimurium* SR5 | 8192 | 1 | 0.125 | 64 |
| *Pseudomonas aeruginosa* PA01 | 512 | 2 | 0.125 | 512 |
| *Pseudomonas aeruginosa* PAER00606 | 8192 | 1 | 0.25 | >8192 |
| *Acinetobacter baumannii* ATCC17978 | 16 | 2 | 0.25 | 256 |
| *Acinetobacter baumannii* CPC35 | 128 | 8 | 8 | 512 |
| *Klebsiella pneumoniae* EH62 | 128 | 1 | 16 | 1024 |

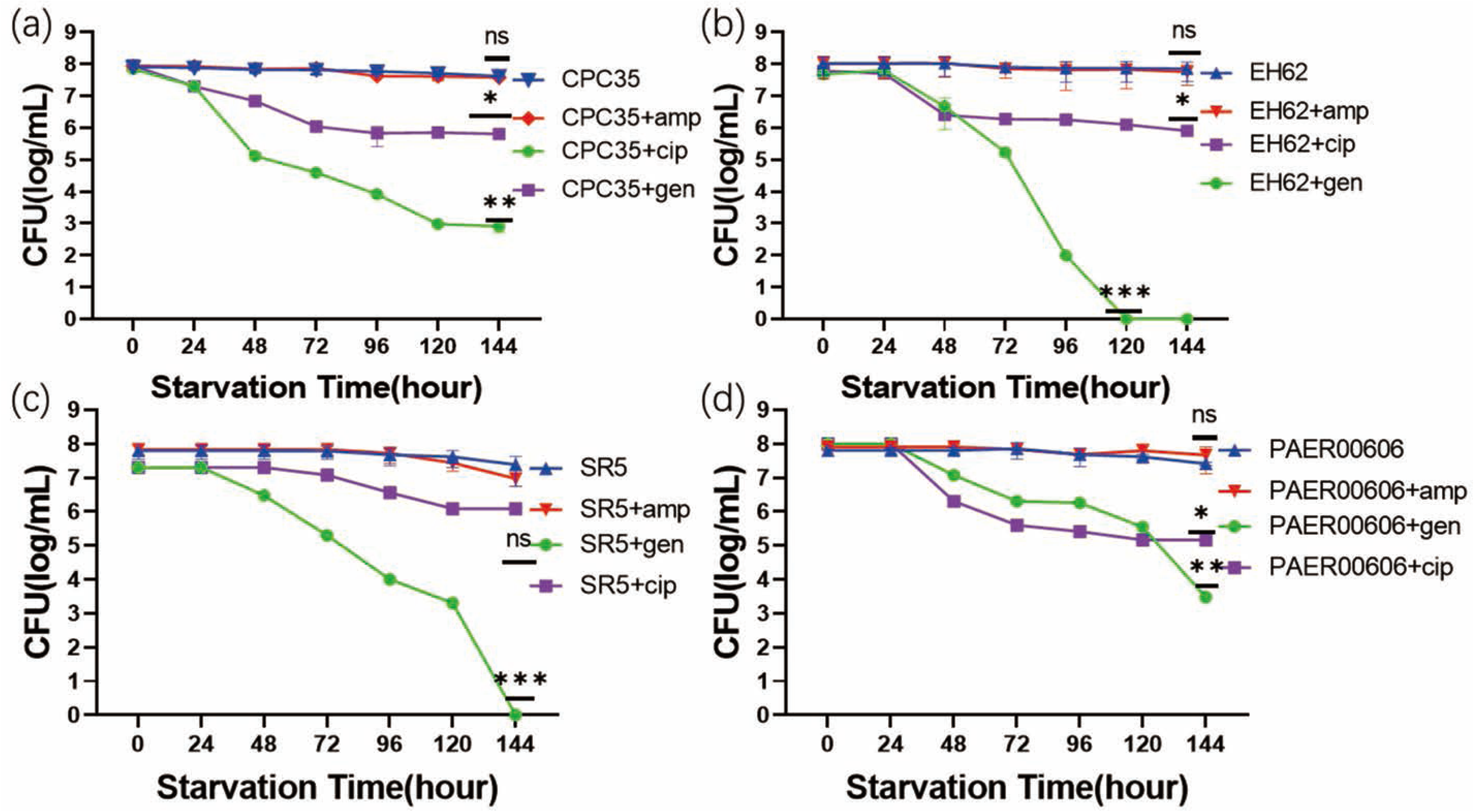

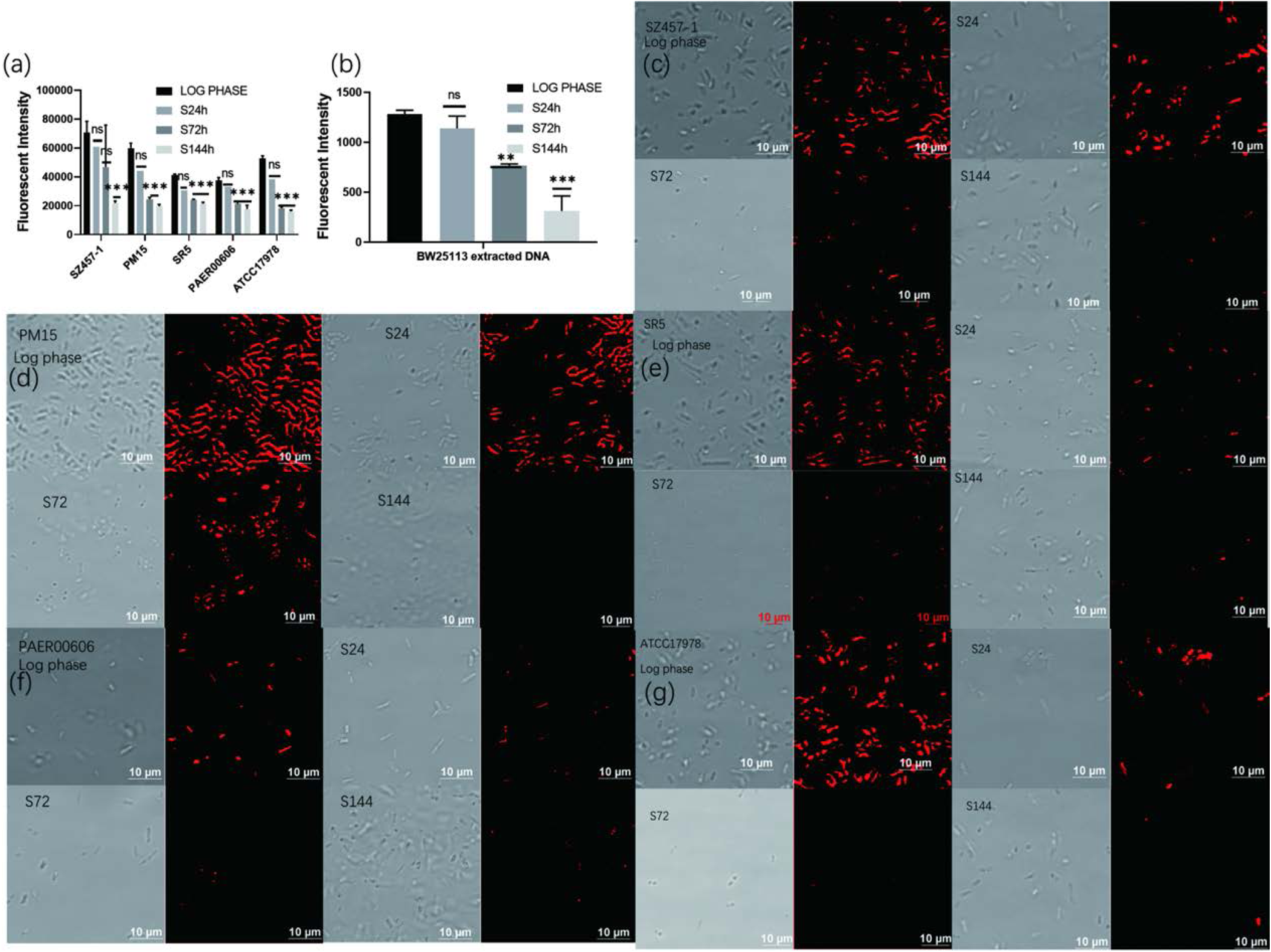

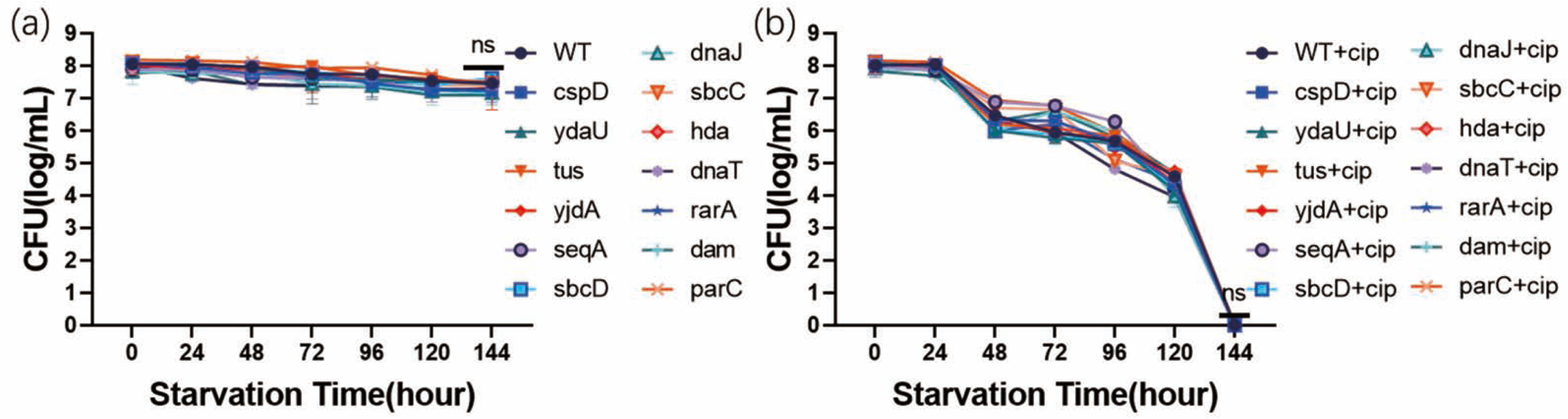

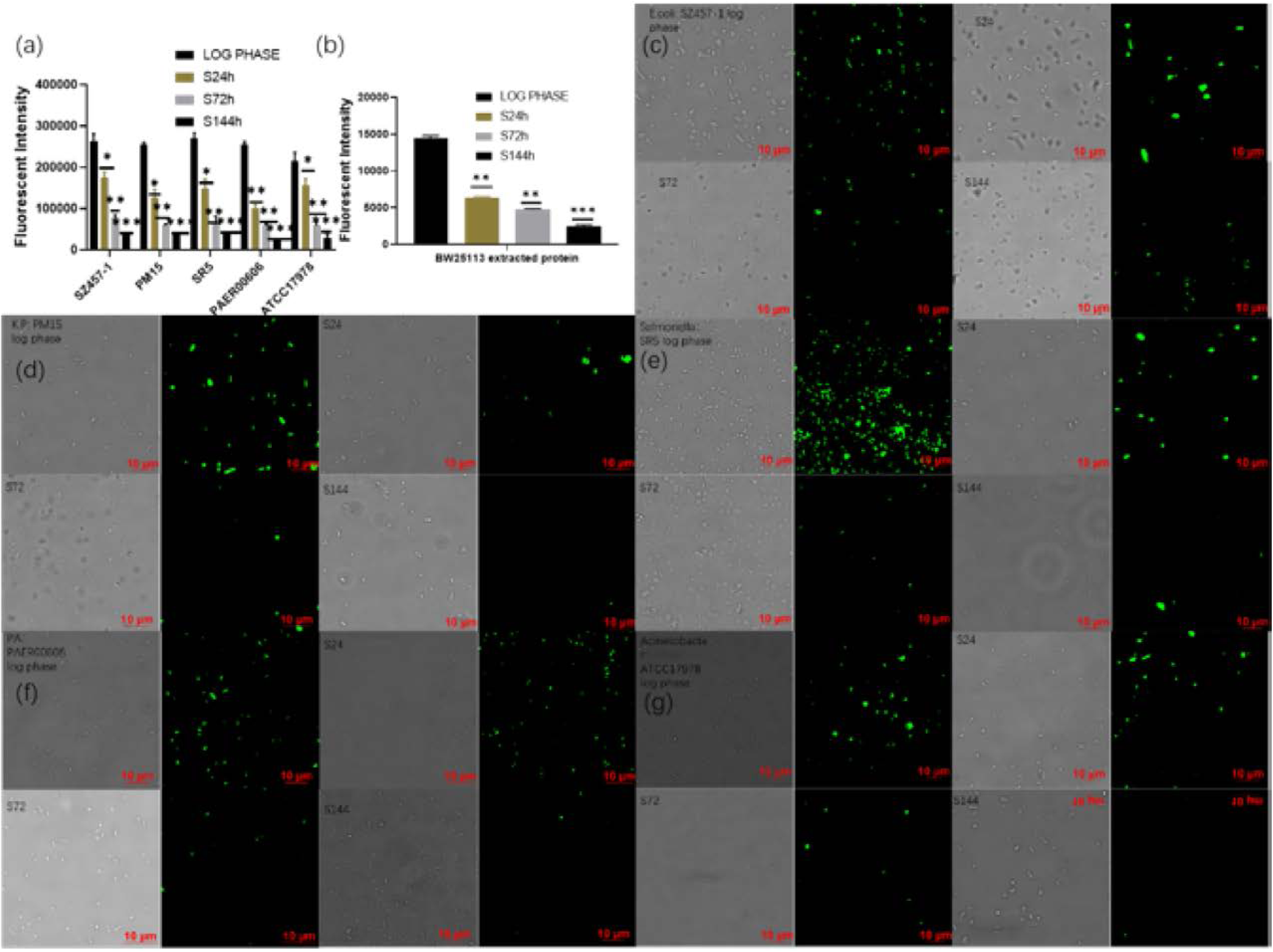

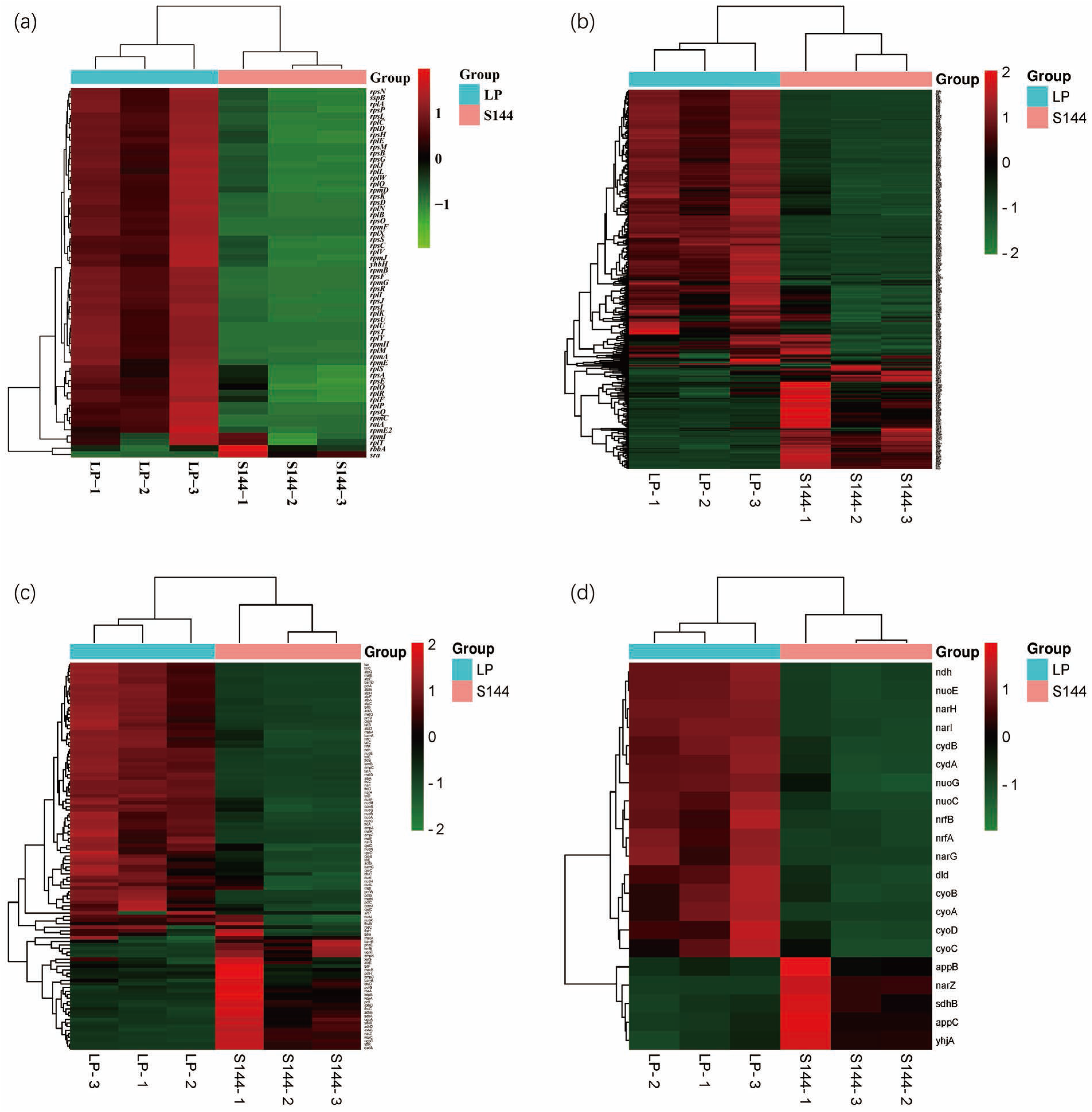

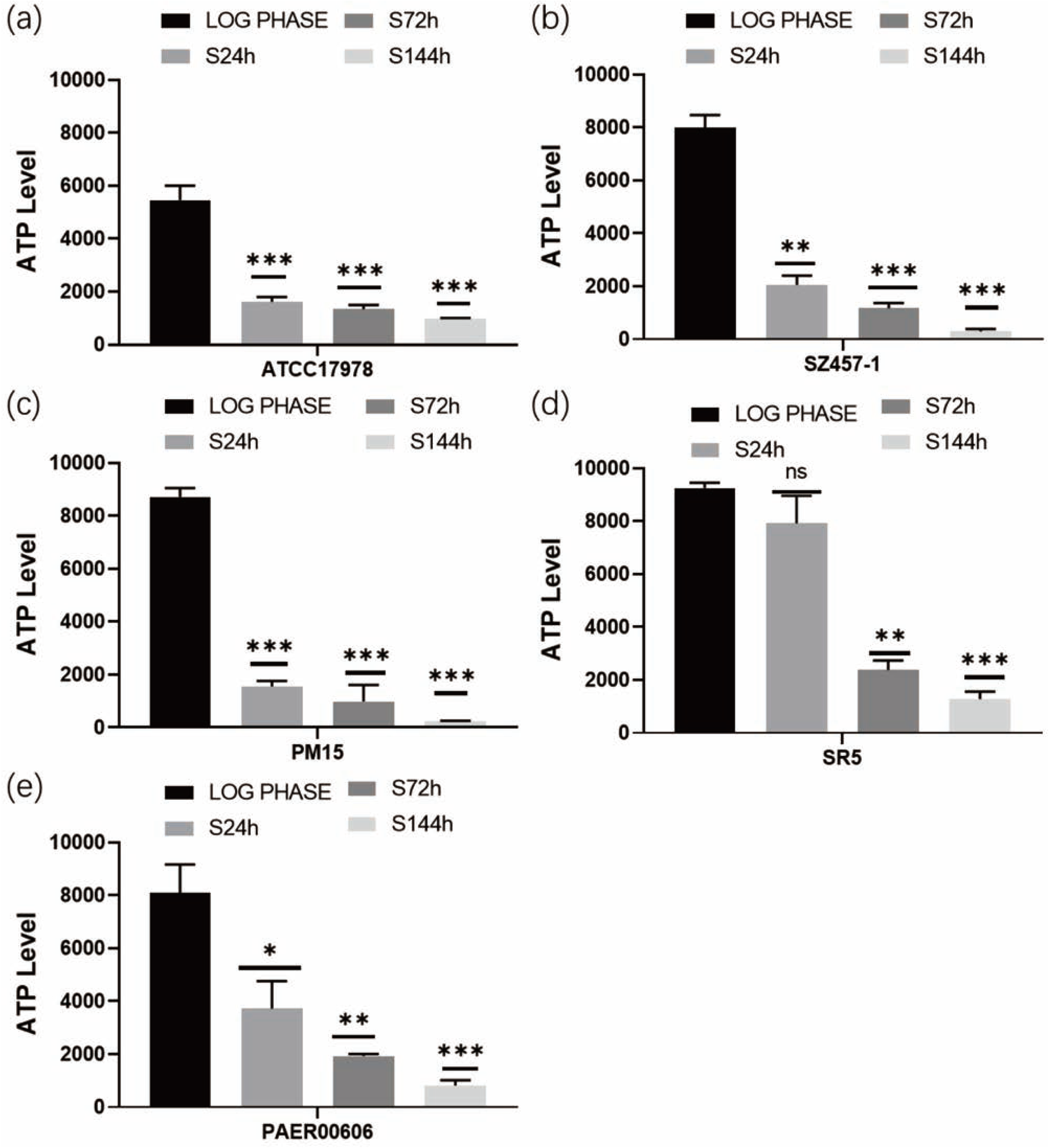

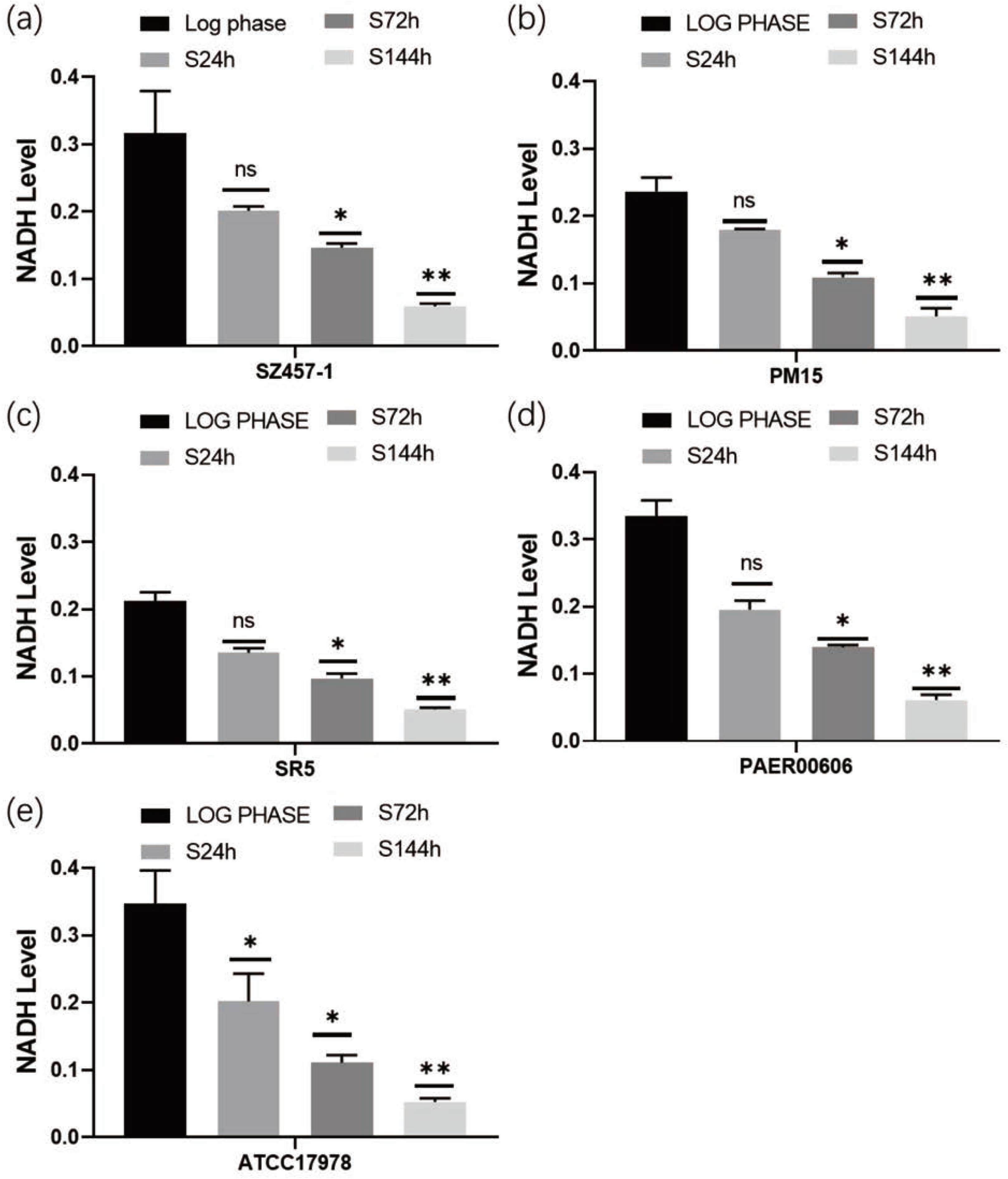

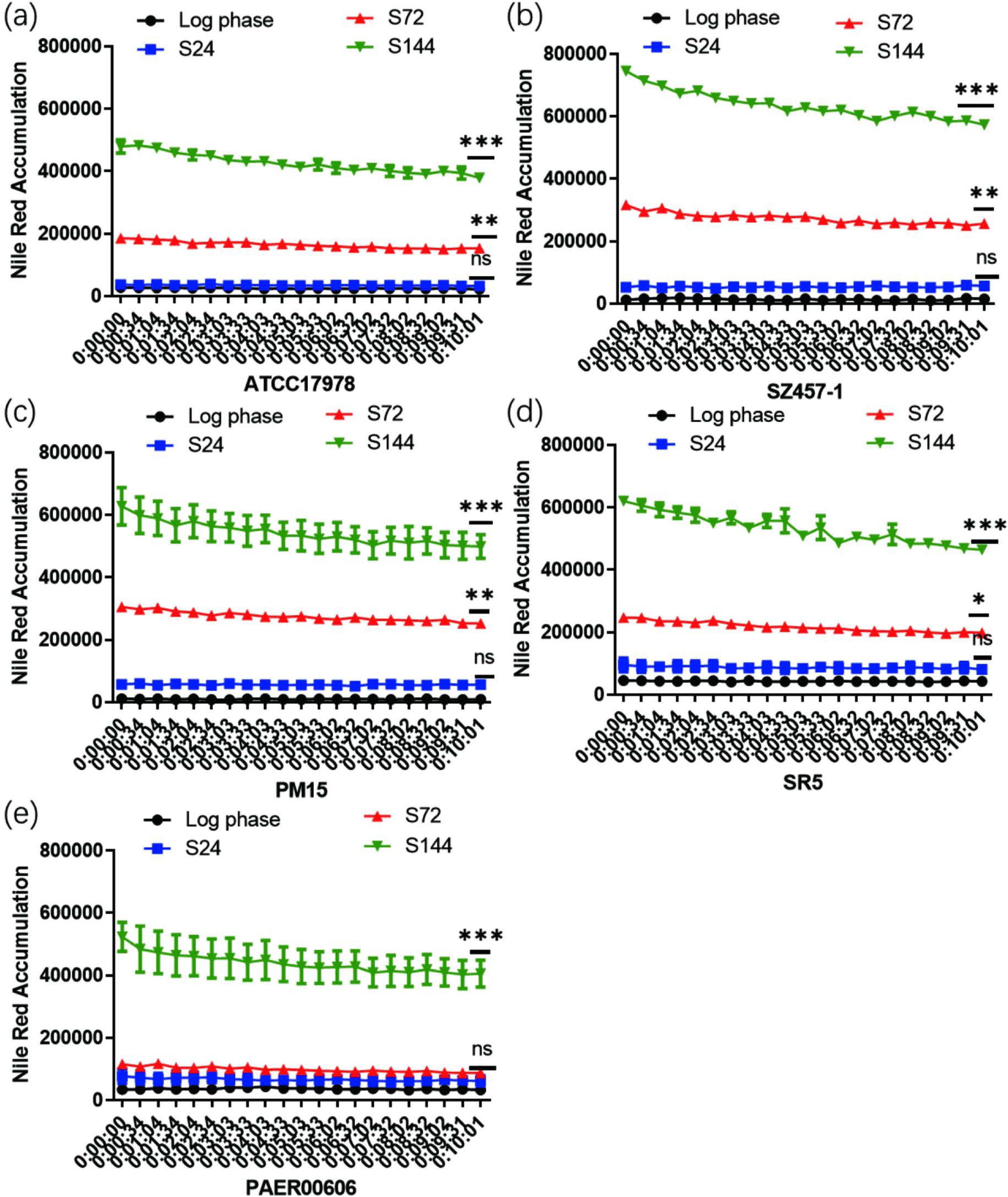

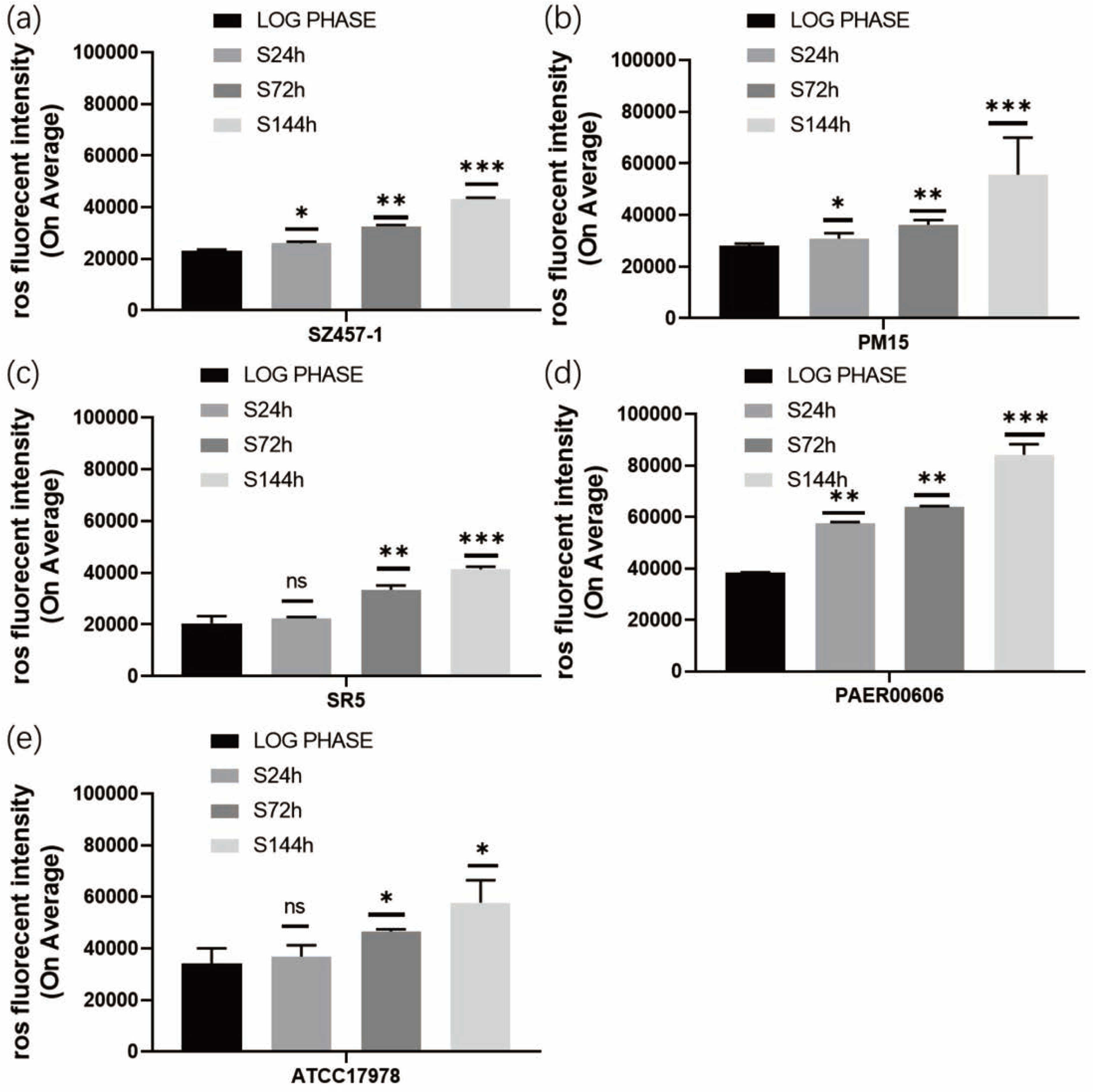

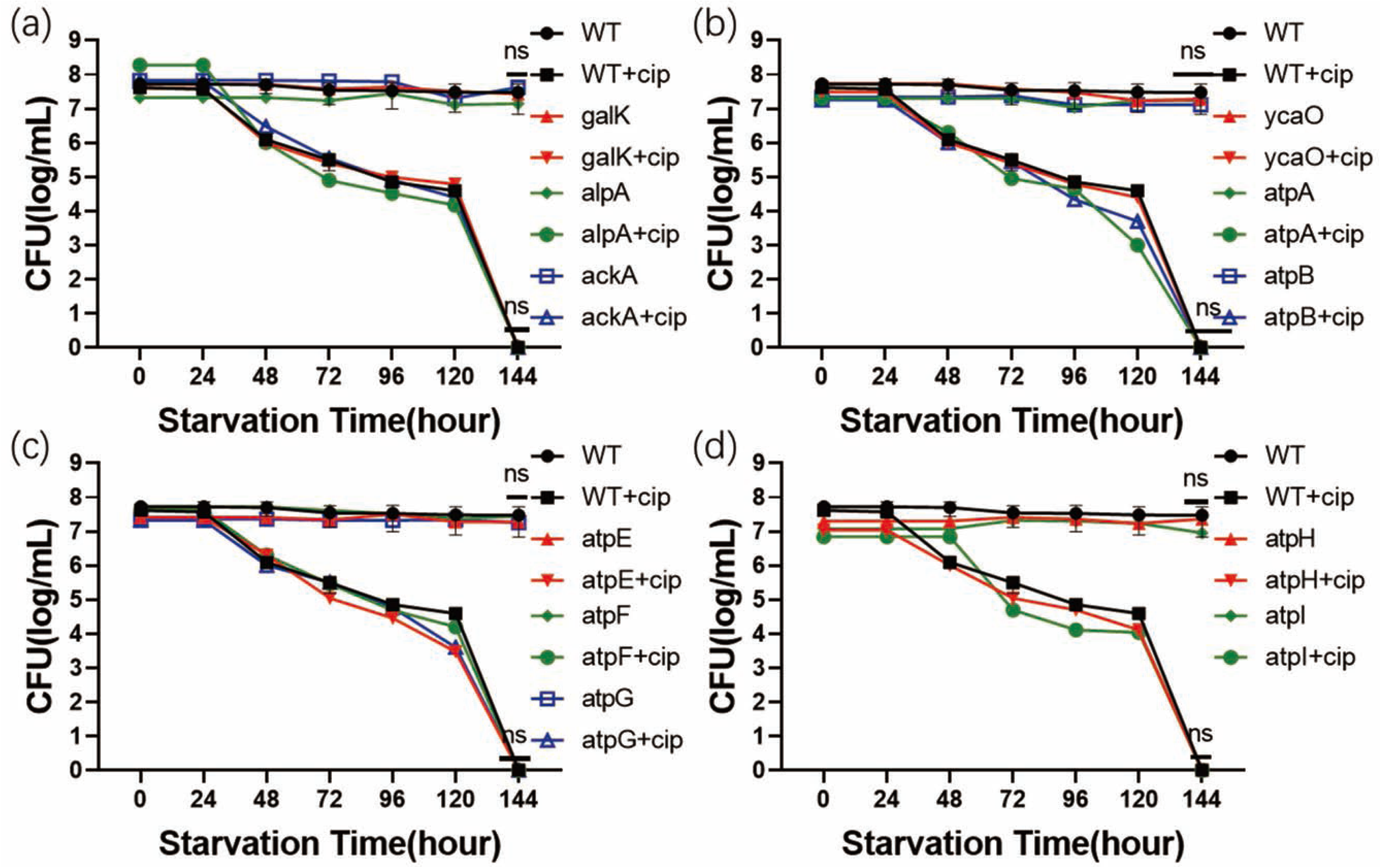

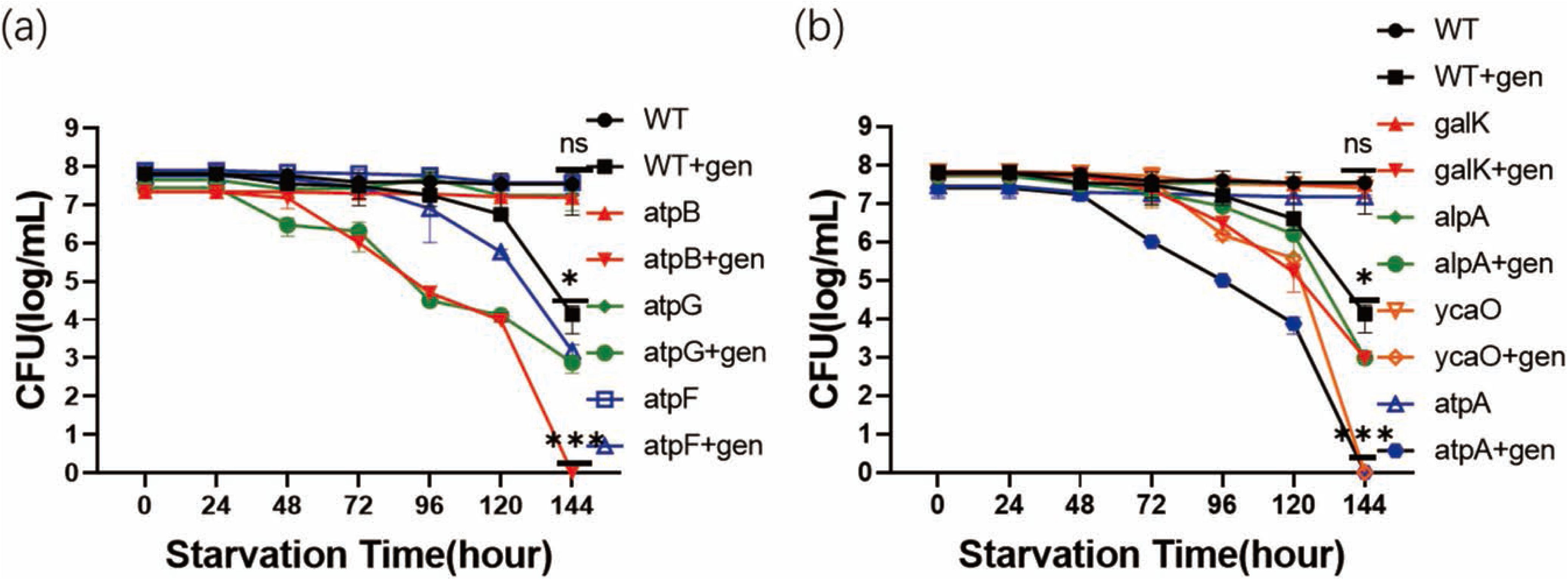

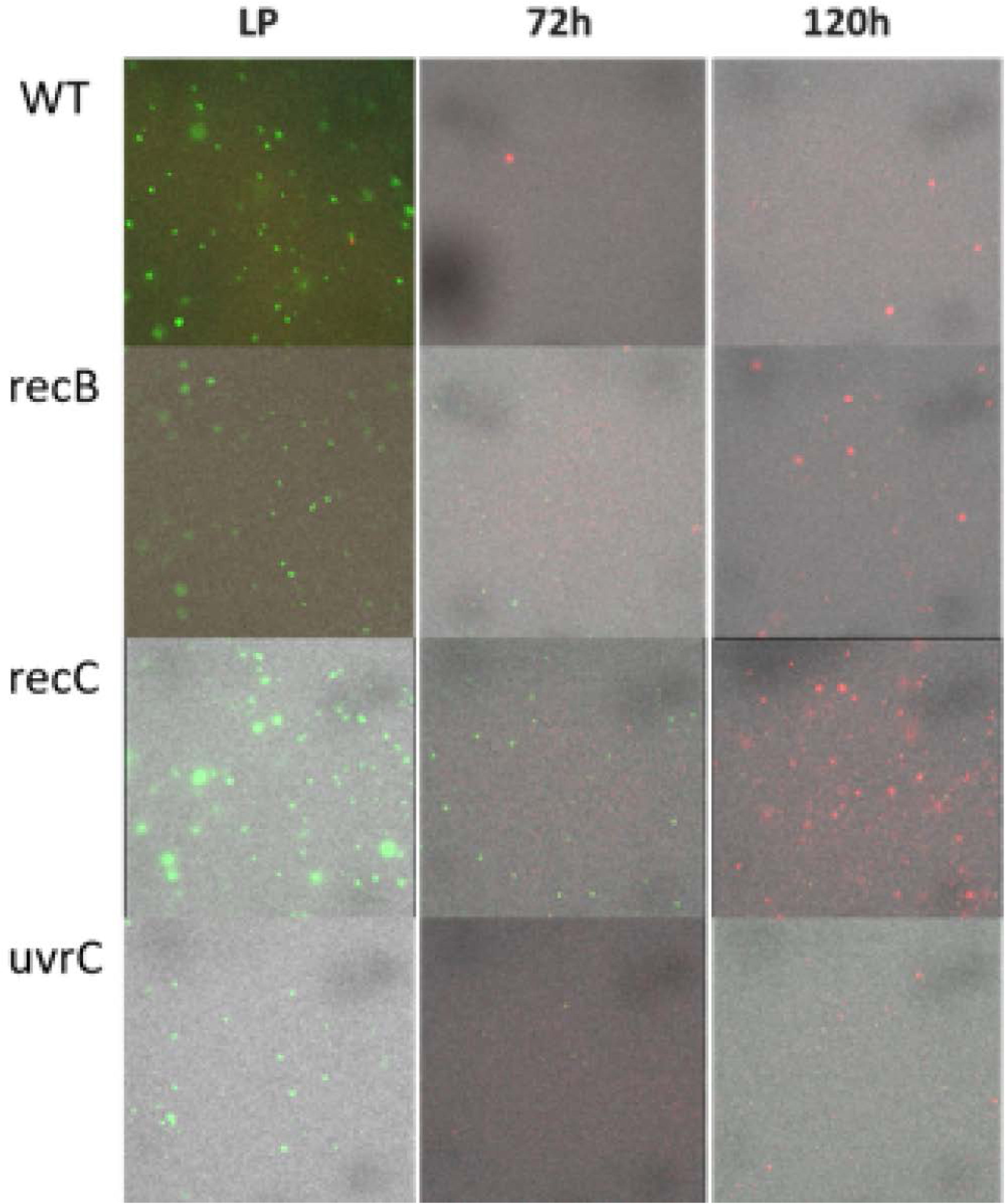

## Reference

Adams, K. N., Takaki, K., Connolly, L. E., Wiedenhoft, H., Winglee, K., Humbert, O., Edelstein, P. H., Cosma, C. L., & Ramakrishnan, L. (2011). Drug tolerance in replicating mycobacteria mediated by a macrophage-induced efflux mechanism. Cell, 145(1), 39–53.

Arnoldini, M., Vizcarra, I. A., Peña-Miller, R., Stocker, N., Diard, M., Vogel, V., Beardmore, R. E., Hardt, W.-D., & Ackermann, M. (2014). Bistable expression of virulence genes in salmonella leads to the formation of an antibiotic-tolerant subpopulation. PLoS biology, 12(8), e1001928.

Bigger, J. (1944). Treatment of Staphyloeoeeal Infections with Penicillin by Intermittent Sterilisation. Lancet, 497–500.

Bokinsky, G., Baidoo, E. E., Akella, S., Burd, H., Weaver, D., Alonso-Gutierrez, J., García-Martín, H., Lee, T. S., & Keasling, J. D. (2013). HipA-triggered growth arrest and β-lactam tolerance in Escherichia coli are mediated by RelA-dependent ppGpp synthesis. Journal of bacteriology, 195(14), 3173–3182.

Cecchini, M. J., Amiri, M., & Dick, F. A. (2012). Analysis of cell cycle position in mammalian cells. JoVE (Journal of Visualized Experiments)(59), e3491.

Chen, S., Cui, S., McDermott, P. F., Zhao, S., White, D. G., Paulsen, I., & Meng, J. (2007). Contribution of target gene mutations and efflux to decreased susceptibility of Salmonella enterica serovar Typhimurium to fluoroquinolones and other antimicrobials. Antimicrobial agents and chemotherapy, 51(2), 535–542.

Fung, D. K., Chan, E. W., Chin, M. L., & Chan, R. C. (2010). Delineation of a bacterial starvation stress response network which can mediate antibiotic tolerance development. Antimicrobial agents and chemotherapy, 54(3), 1082–1093.

Goormaghtigh, F., & Van Melderen, L. (2019). Single-cell imaging and characterization of Escherichia coli persister cells to ofloxacin in exponential cultures. Science advances, 5(6), eaav9462.

Grenier, F., Matteau, D., Baby, V., & Rodrigue, S. (2014). Complete genome sequence of Escherichia coli BW25113. Genome announcements, 2(5), 10.1128/genomea.01038-01014.

Harms, A., Maisonneuve, E., & Gerdes, K. (2016). Mechanisms of bacterial persistence during stress and antibiotic exposure. Science, 354(6318), Article aaf4268. 10.1126/science.aaf4268

Ma, C., Sim, S., Shi, W., Du, L., Xing, D., & Zhang, Y. (2010). Energy production genes sucB and ubiF are involved in persister survival and tolerance to multiple antibiotics and stresses in Escherichia coli. FEMS microbiology letters, 303(1), 33–40.

Mayer, C., Muras, A., Romero, M., López, M., Tomás, M., & Otero, A. (2018). Multiple quorum quenching enzymes are active in the nosocomial pathogen Acinetobacter baumannii ATCC17978. Frontiers in cellular and infection microbiology, 8, 310.

Moosdeen, F., & Cheong, Y. (1989). Enzyme of β-lactam resistant salmonella strains. Journal of Antimicrobial Chemotherapy, 23(5), 797–798.

Orman, M. A., & Brynildsen, M. P. (2015). Inhibition of stationary phase respiration impairs persister formation in E. coli. Nature communications, 6(1), 7983.

Pu, Y., Zhao, Z., Li, Y., Zou, J., Ma, Q., Zhao, Y., Ke, Y., Zhu, Y., Chen, H., & Baker, M. A. (2016). Enhanced efflux activity facilitates drug tolerance in dormant bacterial cells. Molecular cell, 62(2), 284–294.

Radzikowski, J. L., Vedelaar, S., Siegel, D., Ortega, Á. D., Schmidt, A., & Heinemann, M. (2016). Bacterial persistence is an active σS stress response to metabolic flux limitation. Molecular systems biology, 12(9), 882.

Stover, C., Pham, X., Erwin, A., Mizoguchi, S., Warrener, P., Hickey, M., Brinkman, F., Hufnagle, W., Kowalik, D., & Lagrou, M. (2000). Complete genome sequence of Pseudomonas aeruginosa PAO1, an opportunistic pathogen. Nature, 406(6799), 959–964.

Te Winkel, J. D., Gray, D. A., Seistrup, K. H., Hamoen, L. W., & Strahl, H. (2016). Analysis of antimicrobial-triggered membrane depolarization using voltage sensitive dyes. Frontiers in cell and developmental biology, 29.

Volzing, K. G., & Brynildsen, M. P. (2015). Stationary-Phase Persisters to Ofloxacin Sustain DNA Damage and Require Repair Systems Only during Recovery. mBio, 6(5), e00731–00715. 10.1128/mBio.00731-15

Wakamoto, Y., Dhar, N., Chait, R., Schneider, K., Signorino-Gelo, F., Leibler, S., & McKinney, J. D. (2013). Dynamic persistence of antibiotic-stressed mycobacteria. Science, 339(6115), 91–95.

Wan, Y., Wai Chi Chan, E., & Chen, S. (2023). Maintenance and generation of proton motive force are both essential for expression of phenotypic antibiotic tolerance in bacteria. Microbiology spectrum, 11(5), e00832–00823.

Wan, Y., Wang, M., Chan, E. W. C., & Chen, S. (2021). Membrane transporters of the major facilitator superfamily are essential for long-term maintenance of phenotypic tolerance to multiple antibiotics in E. coli. Microbiology Spectrum, 9(3), e01846–01821.

Wang, M. M., Chan, E. W. C., Wan, Y. K., Wong, M. H. Y., & Chen, S. (2021). Active maintenance of proton motive force mediates starvation-induced bacterial antibiotic tolerance in *Escherichia coli*. Communications biology, 4(1), Article 1068. 10.1038/s42003-021-02612-1

Wayne, P. (2011). Clinical and laboratory standards institute. Performance standards for antimicrobial susceptibility testing.

Wood, T. K., Knabel, S. J., & Kwan, B. W. (2013). Bacterial persister cell formation and dormancy. Applied and environmental microbiology, 79(23), 7116–7121.

Zheng, E. J., Andrews, I. W., Grote, A. T., Manson, A. L., Alcantar, M. A., Earl, A. M., & Collins, J. J. (2022). Modulating the evolutionary trajectory of tolerance using antibiotics with different metabolic dependencies. Nature Communications, 13(1), Article 2525. 10.1038/s41467-022-30272-0

